# 7,8-Dihydroxyflavone is a direct inhibitor of pyridoxal phosphatase

**DOI:** 10.1101/2023.10.04.560852

**Authors:** Marian Brenner, Christoph Zink, Linda Witzinger, Angelika Keller, Kerstin Hadamek, Sebastian Bothe, Martin Neuenschwander, Carmen Villmann, Jens Peter von Kries, Hermann Schindelin, Elisabeth Jeanclos, Antje Gohla

## Abstract

Vitamin B6 deficiency has been linked to cognitive impairment in human brain disorders for decades. Still, the molecular mechanisms linking vitamin B6 to these pathologies remain poorly understood, and whether vitamin B6 supplementation improves cognition is unclear as well. Pyridoxal phosphatase (PDXP), an enzyme that controls levels of pyridoxal 5’-phosphate (PLP), the co-enzymatically active form of vitamin B6, may represent an alternative therapeutic entry point into vitamin B6-associated pathologies. However, pharmacological PDXP inhibitors to test this concept are lacking. We now identify a PDXP and age-dependent decline of PLP levels in the murine hippocampus that provides a rationale for the development of PDXP inhibitors. Using a combination of small molecule screening, protein crystallography and biolayer interferometry, we discover, visualize and analyze 7,8-dihydroxyflavone (7,8-DHF) as a direct and potent PDXP inhibitor. 7,8-DHF binds and reversibly inhibits PDXP with low micromolar affinity and sub-micromolar potency. In mouse hippocampal neurons, 7,8-DHF increases PLP in a PDXP-dependent manner. These findings validate PDXP as a druggable target. Of note, 7,8-DHF is a well-studied molecule in brain disorder models, although its mechanism of action is actively debated. Our discovery of 7,8-DHF as a PDXP inhibitor offers novel mechanistic insights into the controversy surrounding 7,8-DHF-mediated effects in the brain.

## INTRODUCTION

Vitamin B6 is an essential micronutrient that plays an important role in the nervous system (1, 2), with the vitamin B6 status affecting cognitive function at any age (3, 4). Population studies indicate that low vitamin B6 levels are common among older people (5), and suggest that vitamin B6 deficiency may influence memory performance and may contribute to age-related cognitive decline (6–9). Vitamin B6 deficiency is also associated with other conditions characterized by impaired learning and memory, including neuropsychiatric disorders (10–12), Alzheimer’s disease (13) and inflammation (14, 15). Nevertheless, the exact molecular mechanisms linking vitamin B6 to these pathologies are often insufficiently understood, and whether vitamin B6 supplementation improves cognition is unclear (4, 5, 16–22).

The term vitamin B6 encompasses the enzymatically interconvertible compounds pyridoxine, pyridoxamine, pyridoxal (collectively referred to as B6 vitamers) and their phosphorylated forms. Among these, only pyridoxal 5′-phosphate (PLP) is co-enzymatically active. In humans, PLP is known to be required for 44 distinct biochemical reactions, including the biosynthesis and/or metabolism of neurotransmitters, amino acids, lipids, and glucose. In addition, B6 vitamers display antioxidant and anti-inflammatory functions (23–26).

Cellular PLP availability in the brain depends on numerous factors, including the intestinal absorption of B6 vitamers, extracellular phosphatases, inter-organ transport and intracellular enzymes and carriers/scavengers involved in PLP formation and homeostasis (2). Specifically, intracellular PLP is formed by the pyridoxal kinase (PDXK)-catalyzed phosphorylation of pyridoxal, or the pyridox(am)ine-5’-phosphate oxidase (PNPO)-catalyzed oxidation of pyridox(am)ine 5’-phosphate to PLP. PLP is highly reactive and can undergo condensation reactions with e.g., primary amino groups or thiol groups in proteins or amino acids. Although the mechanisms of PLP delivery within the cells are still largely unknown, it is clear that the intracellular availability of PLP for co-enzymatic functions depends on PLP carriers/scavengers and on the hydrolytic activity of pyridoxal 5’-phosphate phosphatase (PDXP) (2, 27–30).

We have previously shown that the genetic knockout of PDXP (PDXP-KO) in mice increases brain PLP levels and improves spatial memory and learning, suggesting that elevated PLP levels can improve cognitive functions in this model (30). We therefore reasoned that a pharmacological inhibition of PDXP may be leveraged to increase intracellular PLP levels and conducted a high-throughput screening campaign to identify small-molecule PDXP modulators. Here, we report the discovery and the structural and cellular validation of 7,8-dihydroxyflavone (7,8-DHF) as a preferential PDXP inhibitor. 7,8-DHF is a well-studied molecule in brain disorder models characterized by impaired cognition, and widely regarded as a tropomyosin receptor kinase B (TrkB) agonist with brain-derived neurotrophic factor (BDNF)-mimetic activity (31). However, a direct TrkB agonistic activity of 7,8-DHF has been called into question (32–36). Our serendipitous discovery of 7,8-DHF as a direct PDXP inhibitor provides an alternative mechanistic explanation for 7,8-DHF-mediated effects. More potent, efficacious, and selective PDXP inhibitors may be useful future tools to explore a possible benefit of elevated PLP levels in brain disorders.

## RESULTS

### PDXP activity controls PLP levels in the hippocampus

The hippocampus is important for age-dependent memory consolidation and learning, and impaired memory and learning is associated with PLP deficiency (3). To study a possible contribution of PDXP and/or PDXK to age-related PLP homeostasis in the hippocampus, we performed Western blot analyses in young versus older mice. Unexpectedly, we found that both PDXP and PDXK expression levels were markedly higher in hippocampi of middle-aged than of juvenile animals (**Fig. 1a**). These data suggest an accelerated hippocampal PLP turnover in older mice, consistent with previous findings in senescent mice (37).

**Figure 1.**
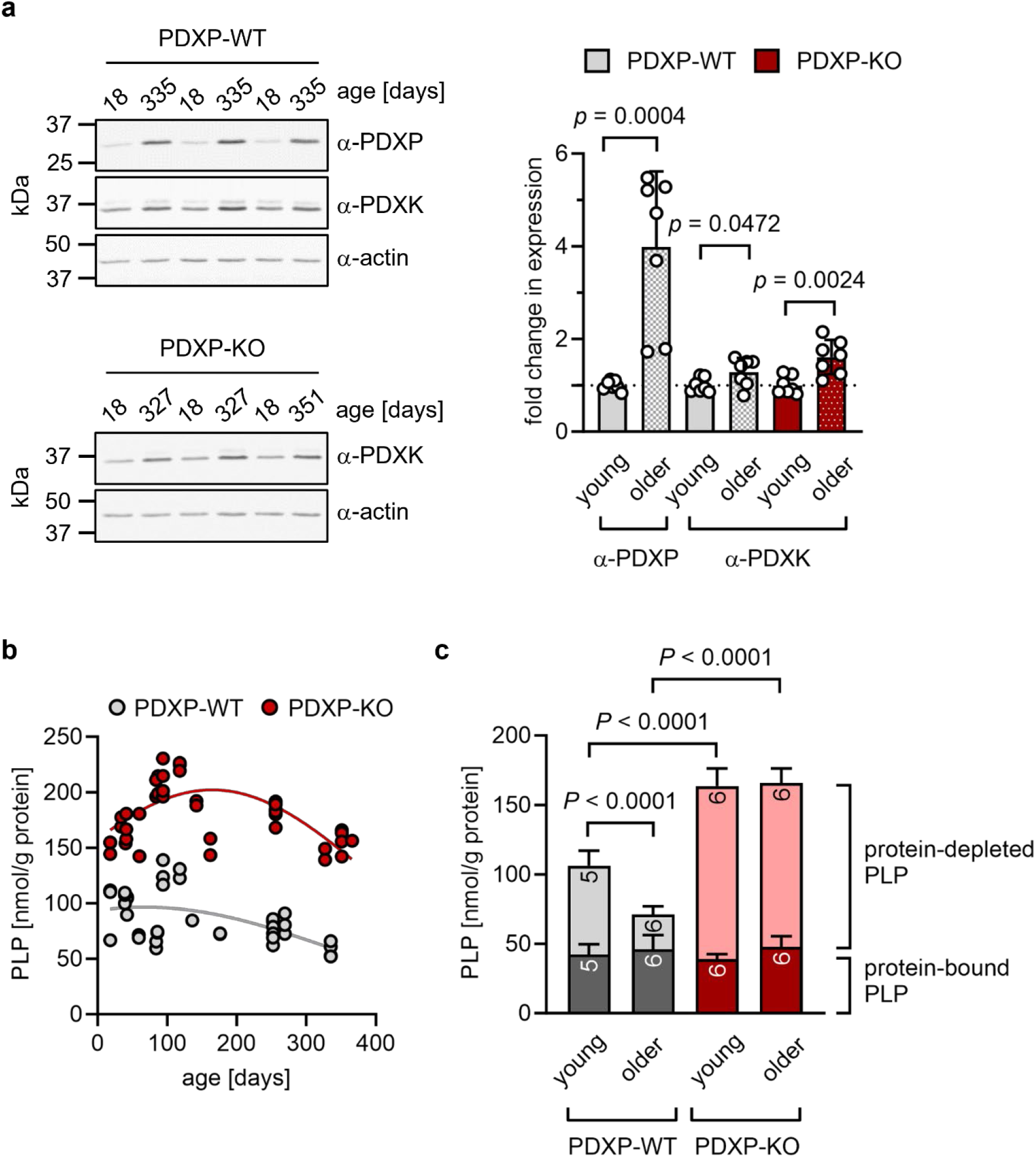
Role of PDXP in hippocampal PLP homeostasis. (**a**) Age-dependent expression of PDXK and PDXP in murine hippocampi. *Left panels,* representative Western blots of three hippocampi for each genotype. The same blots were reprobed with α-actin antibodies as a loading control. The age of the investigated mice is indicated above the blots. *Right panel,* densitometric quantification of hippocampal PDXP and PDXK Western blot signals, corrected by the corresponding actin signals. Young mice were 18-42 days old, older mice were 252-351 days old; *n*=7 hippocampi were analyzed per group. Data are mean values ± S.D. Statistical analysis was performed with unpaired, two-sided *t*-tests; *p*-values are indicated. (**b**) Age-dependent, total PLP-concentrations in isolated hippocampi of PDXP-WT and PDXP-KO mice. PLP was derivatized with semicarbazide and analyzed by HPLC. Each symbol represents the result of the PLP determination in an individual hippocampus. Data were fitted by Gaussian least-squares analyses. (**c**) Determination of protein-bound and protein-depleted PLP in PDXP-WT and PDXP-KO hippocampal lysates of young (18-42 days old) and older mice (252-352 days old). The number of analyzed hippocampi is indicated in the bars. Data are mean values ± S.D. Statistical analysis was performed with two-way ANOVA and Tukey’s multiple comparisons test. Significant differences (adjusted *P*-values) in protein-depleted PLP levels are indicated. The exact age of analyzed mice is listed in **Figure 1 – supplementary figure 1**. **Source data** are available for this Figure.

An analysis of total hippocampal PLP levels in PDXP-WT and PDXP-KO mice showed an age-dependent profile. PLP levels appeared to peak around 3 months of age (possibly reflecting PLP-dependent neurotransmitter biosynthesis and metabolism during the postnatal developmental period) and descended back to juvenile levels by 12 months of age in both genotypes. Although total hippocampal PLP levels in PDXP-KO mice also decreased with age, they consistently remained above PLP levels in control mice (**Fig. 1b**; two-tailed, unpaired *t*-test of PLP levels in PDXP-WT vs. PDXP-KO hippocampi, all ages combined: *p*<0.0001).

PLP is protected from hydrolysis by binding to proteins, and PDXP is expected to dephosphorylate only non-protein-bound PLP (38). To test this, we prepared protein-depleted PLP fractions from PDXP-WT and PDXP-KO hippocampal lysates using 3 kDa molecular weight cutoff centrifugal filters. The quantification of PLP in these fractions demonstrated that PDXP loss indeed only increased the pool of protein-depleted PLP, both in young (18-42 days old) and older mice (252-352 days old, corresponding to mature/middle-aged mice), whereas the levels of protein-bound PLP remained unchanged (**Fig. 1c**). While the hippocampal levels of non-protein-bound PLP dropped by about 60% over this time span in PDXP-WT mice, they remained elevated in PDXP-KO mice (∼2-fold higher in younger, and ∼5-fold higher in older PDXP-KO compared to the respective PDXP-WT; see **Fig. 1 – figure supplement 1** for exact mouse ages). We conclude that hippocampi of older mice are characterized by a specific decrease in the levels of non-protein-bound PLP, and that this age-dependent PLP loss is dependent on PDXP activity. These observations establish that PDXP is a critical determinant of PLP levels in the murine hippocampus and suggest that intracellular PLP deficiency may be alleviated by PDXP inhibition.

### A high-throughput screening campaign identifies 7,8-dihydroxyflavone as a PDXP inhibitor

Pharmacological small-molecule PDXP inhibitors are currently lacking. To identify PDXP inhibitor candidates, we screened the FMP small molecule repository containing 41,182 compounds for molecules able to modulate the phosphatase activity of recombinant, highly purified murine PDXP (see **Fig. 2 – figure supplement 1** for a schematic of the screening campaign). Difluoro-4-methylumbelliferyl phosphate (DiFMUP) was used as a fluorogenic phosphatase substrate in a primary screen. Compounds that altered DiFMUP fluorescence by ≥50% (activator candidates) or ≤25% (inhibitor candidates) were subjected to EC_50_/IC_50_ value determinations. Of these, 46 inhibitor hits were selected and counter-screened against phosphoglycolate phosphatase (PGP), the closest PDXP relative (39, 40). Eleven of the PDXP inhibitor hits (with an IC_50_ PDXP <20 µM, and IC_50_ PDXP < IC_50_ PGP or no activity against PGP) were subsequently validated in a secondary assay, using PLP as a physiological PDXP substrate (see **Fig. 2 – figure supplement 2** for all 11 inhibitor hits). Only one PDXP-selective inhibitor hit (7,8-dihydroxyflavone/7,8-DHF, a naturally occurring flavone) blocked PDXP-catalyzed PLP dephosphorylation (IC_50_ ∼1 µM).

In vitro activity assays using PLP as a substrate confirmed that 7,8-DHF directly blocks murine and human PDXP activity with submicromolar potency and an apparent efficacy of ∼50% (**Fig. 2a, b**). We next examined whether commercially available 7,8-DHF analogs might be more potent or efficacious PDXP inhibitors. We tested flavone, 3,7-dihydroxyflavone, 5,7-dihydroxyflavone (also known as chrysin), 3,5,7-trihydroxyflavon (galangin), 5,6,7-trihydroxyflavone (baicalein) and 3,7,8,4’-tetrahydroxyflavone. **Figure 2b** shows that of the tested 7,8-DHF analogs, only 3,7,8,4’-tetrahydroxyflavone was able to inhibit PDXP, albeit with an IC_50_ of 2.5 µM and thus slightly less potently than 7,8-DHF. These results suggest that hydroxyl groups in positions 7 and 8 of the flavone scaffold are required for PDXP inhibition. The efficacy of PDXP inhibition by 3,7,8,4’-tetrahydroxyflavone was not substantially increased at concentrations >40 µM (relative PDXP activity at 40 µM: 0.46 ± 0.05; at 70 µM: 0.38 ± 0.15; at 100 µM: 0.37 ± 0.09; data are mean values ± S.D. of *n*=6 experiments). Concentrations >100 µM could not be assessed due to impaired PDXP activity at the DMSO concentrations required for solubilizing the flavone.

**Figure 2:**
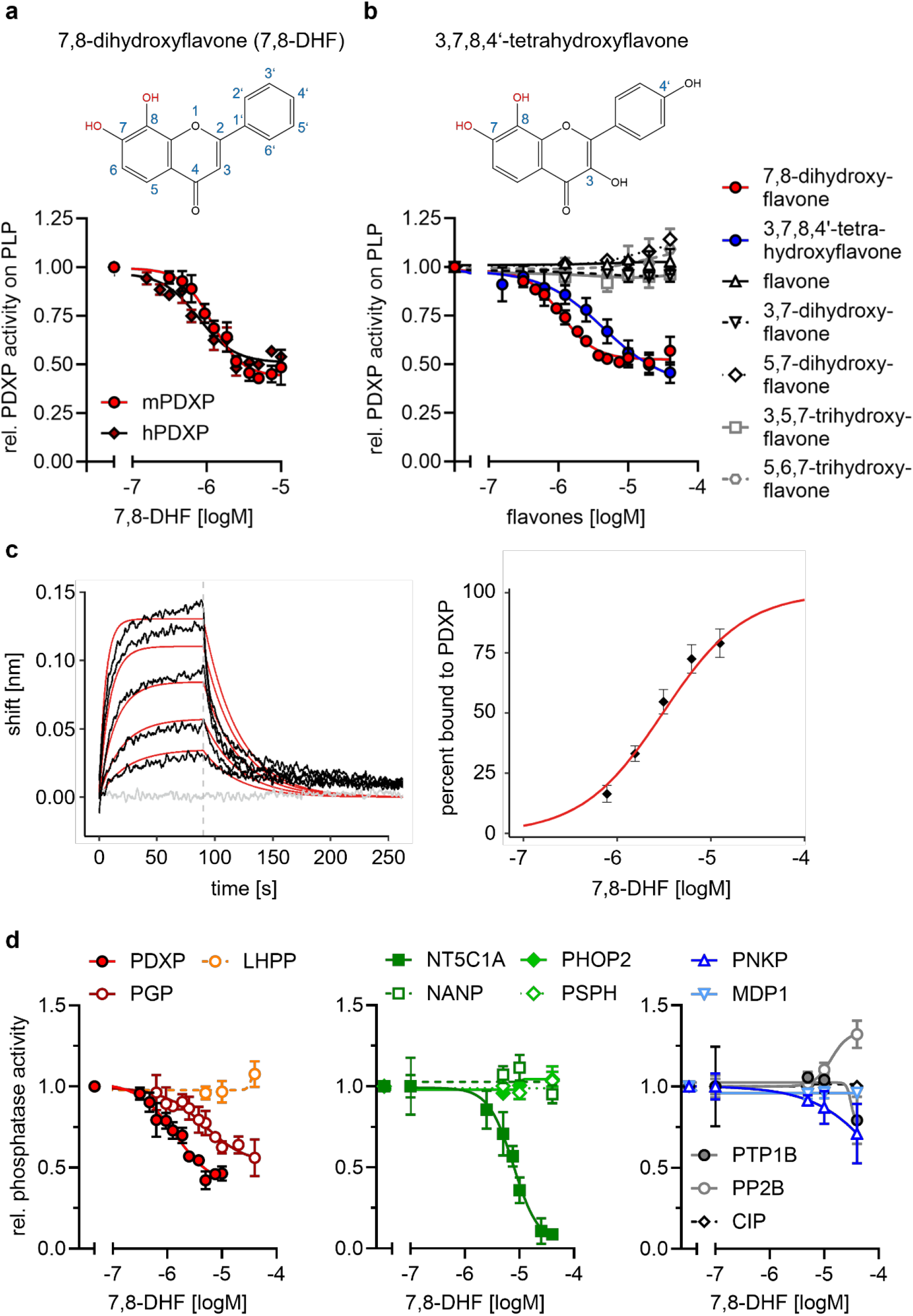
Characterization of the 7,8-DHF/PDXP interaction. (**a**) Determination of half-maximal inhibitory constants (IC_50_) of 7,8-DHF (2D-structure shown on top) for purified murine or human PDXP, using pyridoxal 5’-phosphate (PLP) as a substrate. Phosphatase activities in the presence of 7,8-DHF were normalized to the respective enzyme activities measured in the presence of the DMSO solvent control. Data are mean values ± S.D. of *n*=3 (human PDXP) and *n*=4 (murine PDXP) independent experiments. (**b**) IC_50_ values of different flavones for purified murine PDXP with PLP as a substrate. Phosphatase activities in the presence of flavones were normalized to the respective enzyme activities in the presence of the DMSO solvent control. All data are mean values ± S.D. The inhibition of PDXP by 3,7,8-trihydroxyflavone-4’-hydroxyphenyl (2D-structure shown on top) was assessed in *n*=6 independent experiments. All other data are from *n*=3 biologically independent experiments. Apparently missing error bars are hidden by the symbols. (**c**) Biolayer interferometry (BLI) measurements of the interaction of 7,8-DHF with purified murine PDXP. *Left panel*, example sensorgram overlayed with the global 1:1 binding model (red) and the negative control (gray). The dashed line indicates the start of the dissociation phase. *Right panel*, steady-state dose-response analysis for 7,8-DHF based on *n*=4 measurements. (**d**) Sensitivity of the indicated phosphatases to 7,8-DHF. Phosphatase activities in the presence of 7,8-DHF were normalized to the respective enzyme activities measured in the presence of the DMSO solvent control. Data are mean values ± S.D. of *n*=4 (PGP) or *n*=3 independent experiments (all other phosphatases). Phosphatase substrates and HAD phosphatase cap types are indicated in parentheses. PDXP, pyridoxal-5’-phosphate phosphatase (pyridoxal 5’-phosphate, C2); PGP, phosphoglycolate phosphatase (2-phosphoglycolate; C2); LHPP, phospholysine phosphohistidine inorganic pyrophosphate phosphatase (imidodiphosphate; C2); NT5C1A, soluble cytosolic 5’-nucleotidase 1A (AMP; C1); NANP, N-acetylneuraminate 9-phosphate phosphatase (6-phosphogluconate; C1); PHOP2, phosphatase orphan 2 (pyridoxal 5’-phosphate; C1); PSPH, phosphoserine phosphatase (O-phospho-L-serine; C1); PNKP, polynucleotide kinase phosphatase (3-phospho-oligonucleotide; C0); MDP1, magnesium-dependent phosphatase-1 (D-ribose-5-phosphate; C0); PTP1B (protein tyrosine phosphatase 1B; EGFR phospho-peptide); PP2B, protein phosphatase 2B/calcineurin (PKA regulatory subunit type II phospho-peptide); CIP, calf intestinal phosphatase (*p*NPP). **Source data** are available for this Figure.

We used a biolayer interferometry (BLI) optical biosensing technique to further characterize the binding of 7,8-DHF to PDXP (**Fig. 2c**). Consistent with a specific interaction, 7,8-DHF binding to PDXP was concentration-dependent and fully reversible. As a result of the poor solubility of the molecule, a saturation of the binding site was not experimentally accessible. Steady-state analysis of a 7,8-DHF serial dilution series yielded an affinity (K_D_) value of 3.1 ± 0.3 µM (data are mean values ± S.E. of *n*=4 measurements; see **Fig. 2 – figure supplement 3** for the three other measurements) using a 1:1 dose-response model. Global analysis of the sensorgrams assuming a 1:1 binding model resulted in an affinity of 2.6 ± 0.5 µM, in line with the steady-state results (**Fig. 2c**). As expected, 5,7-dihydroxyflavone showed no signal in the BLI, in line with previous experiments (see **Fig. 2b**). With its molecular size of 254 Da and its physicochemical properties, 7,8-DHF is a typical fragment-like molecule (41). Typical association rate constants (k_on_) for fragments are limited by the rate of diffusion and are higher than 10^6^·M^−1^s^−1^. Interestingly, 7,8-DHF showed a slow k_on_ of 1.05·10^4^ M^−1^s^−1^, which is atypical and rarely found for fragment-like molecules (42), and a k_off_ rate of 0.03 s^−1^. With the commonly used estimation of ΔG ∼ pK_D_ and a heavy atom number of 19, 7,8-DHF shows a high ligand efficiency of 0.39, which makes it an interesting molecule for further medicinal chemistry optimization. Taken together, these data support a direct and reversible physical interaction between 7,8-DHF and PDXP that leads to PDXP inhibition.

### Selectivity of 7,8-DHF

PDXP is a member of the large family of haloacid dehalogenase (HAD)-type hydrolases (43). HAD phosphatases are Mg^2+^-dependent phospho-aspartate transferases that consist of a Rossman-like catalytic core linked to a cap domain. The insertion site, structure and size of the cap define the substrate selectivity of the respective enzyme. The “capless” C0-type HAD phosphatases contain either a very small or no cap, resulting in an accessible catalytic cleft that enables the dephosphorylation of macromolecular substrates. Larger C1 or C2 caps act as a roof for the entrance to the active site; most C1/C2-capped HAD phosphatases consequently dephosphorylate small molecules that can gain access to the catalytic cleft. Cap domains also contain so-called substrate specificity loops that contribute to substrate coordination. Hence, caps are distinguishing features of HAD phosphatases (38, 43–45).

To probe the selectivity of 7,8-DHF for PDXP, a C2-capped HAD phosphatase, we tested eight other mammalian HAD phosphatases, including two other C2-, four C1- and two C0-type enzymes. In addition, we analyzed the activity of 7,8-DHF towards the prototypical tyrosine phosphatase PTP1B (which is known to be sensitive to specific flavonoids, ref. (46)); the serine/threonine protein phosphatase calcineurin (PP2B); and a DNA/RNA-directed alkaline phosphatase (calf intestinal phosphatase, CIP) (**Fig. 2d**). When assayed at nominal concentrations of 5, 10 and 40 µM (i.e., up to ∼40-fold above the IC_50_ value for PDXP-catalyzed PLP-dephosphorylation), 7,8-DHF was completely inactive against six of the tested enzymes. At the highest tested concentration of 40 µM, 7,8-DHF weakly inhibited PTP1B and the polynucleotide kinase-3’-phosphatase PNKP and appeared to increase the activity of calcineurin. As expected, 7,8-DHF inhibited PGP, the closest PDXP relative, with an IC_50_ value of 4.8 µM. This result is consistent with the criteria applied during the initial counter-screen (see above). In addition to PGP, 7,8-DHF inhibited the C1-capped soluble cytosolic 5’-nucleotidase 1A (NT5C1A) with an IC_50_ value of ∼10 µM. NT5C1A is an AMP hydrolase expressed in skeletal muscle and heart (47), which is also sensitive to inhibition by small molecules that target the closest PDXP-relative PGP (40). Together, the selectivity analysis of 7,8-DHF on a total of 12 structurally and functionally diverse protein- and non-protein-directed phosphatases show that 7,8-DHF preferentially inhibits PDXP, and that higher 7,8-DHF concentrations can also target the PDXP paralog PGP and the nucleotidase NT5C1A.

### Mode of PDXP inhibition

To probe the mechanism of PDXP inhibition, we assayed the steady state kinetics of PLP dephosphorylation in the presence of increasing 7,8-DHF concentrations (**Table 1**). Analysis of the derived kinetic constants demonstrated that 7,8-DHF increased the K_M_ up to ∼2-fold, and slightly reduced v_max_ values ∼0.7-fold. Thus, 7,8-DHF mainly exhibits a mixed mode of PDXP inhibition, which is predominantly competitive.

**Table 1.** Kinetic constants of PDXP-catalyzed PLP hydrolysis in the presence of 7,8-DHF.

| 7,8-DHF [$\mu\text{M}$] | 0 | 1.0 | 1.5 | 2.0 | 3.0 | 5.0 | 10.0 |
| --- | --- | --- | --- | --- | --- | --- | --- |
| $K_M$ [$\mu\text{M}$] | 14.98<br>$\pm 1.28$ | 18.54<br>$\pm 6.24$ | 20.20<br>$\pm 6.19$ | 18.97<br>$\pm 5.15$ | 24.83<br>$\pm 2.61$ | 32.96<br>$\pm 2.13$ | 30.61<br>$\pm 2.57$ |
| $v_{\text{max}}$<br>[$\mu\text{mol}/\text{min}/\text{mg}$] | 1.08<br>$\pm 0.04$ | 0.95<br>$\pm 0.01$ | 0.89<br>$\pm 0.04$ | 0.85<br>$\pm 0.02$ | 0.80<br>$\pm 0.04$ | 0.81<br>$\pm 0.05$ | 0.74<br>$\pm 0.06$ |
| $k_{\text{cat}}$ [$\text{s}^{-1}$] | 0.57<br>$\pm 0.02$ | 0.5<br>$\pm 0.01$ | 0.47<br>$\pm 0.02$ | 0.45<br>$\pm 0.01$ | 0.42<br>$\pm 0.02$ | 0.43<br>$\pm 0.03$ | 0.39<br>$\pm 0.03$ |
| $k_{\text{cat}}/K_M$ [$\text{s}^{-1}\cdot\text{M}^{-1}$]<br>($\times 10^{-4}$) | 3.93<br>$\pm 0.29$ | 3.27<br>$\pm 0.84$ | 2.75<br>$\pm 0.67$ | 2.72<br>$\pm 0.66$ | 1.72<br>$\pm 0.10$ | 1.31<br>$\pm 0.06$ | 1.29<br>$\pm 0.01$ |

The data are mean values ± S.E.M. of $n=3$ independent experiments, except for the solvent control samples ( $n=6$ ). Curves were fitted and parameters $K_M$ (Michaelis–Menten constant); $v_{\text{max}}$ , (maximum enzyme velocity); $k_{\text{cat}}$ (turnover number) were derived using the Michaelis–Menten model in GraphPad Prism 9.5.1. The $k_{\text{cat}}$ values were calculated from the maximum enzyme velocities using a molecular mass of 31,828 Da for PDXP. DMSO concentrations were kept constant (0.1% DMSO under all conditions, including the solvent control samples). **Source data** are available for this Table.

### Co-crystal structures of PDXP bound to 7,8-DHF

To investigate the mechanism of PDXP inhibition in more detail, we co-crystallized homo-dimeric, full-length murine and human PDXP (mPDXP, hPDXP) with this compound. 7,8-DHF-bound murine PDXP co-crystallized with phosphate in the cubic space group I23, with protomer A containing the inhibitor and protomer B representing an inhibitor-free state (**Fig. 3 – figure supplement 1a**). The structure was refined following molecular replacement with full-length murine PDXP (here referred to as apo-mPDXP; Protein Data Bank/PDB entry 4BX3) to a resolution of 2.0 Å resulting in an R_work_ of 18.4% and an R_free_ of 21.1% (PDB code 8QFW). We additionally obtained two co-crystal structures of human PDXP with 7,8-DHF; one in a phosphate-containing, and one in a phosphate-free form. Both forms crystallized in the tetragonal space group P 4_3_2_1_2, and each protomer of both structures contained the inhibitor (**Fig. 3a**). These structures were refined following molecular replacement with full-length human PDXP (PDB entry 2P27, here referred to as apo-hPDXP) to a resolution of 1.5 Å resulting in an R_work_/R_free_ of 17.0/19.4% (phosphate-bound 7,8-DHF-hPDXP, PDB code 9EM1), and to a resolution of 1.5 Å resulting in an R_work_/R_free_ of 18.2/20.5% (phosphate-free 7,8-DHF-hPDXP, PDB code 8S8A). Data collection and refinement statistics are summarized in **Table 2**.

**Figure 3.**
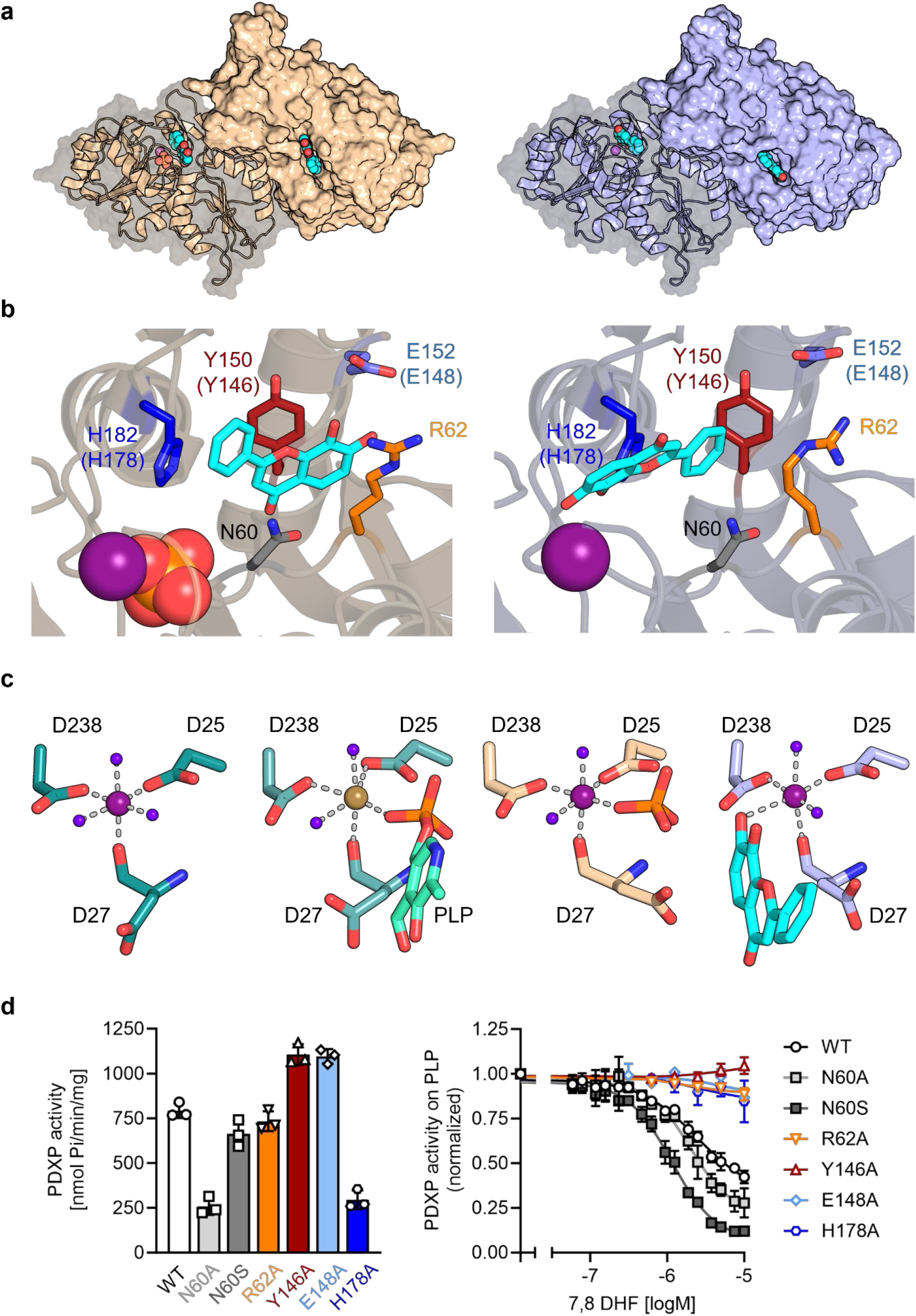
X-ray crystal structures of human PDXP in complex with 7,8-DHF. (**a**) The models were refined to a resolution of 1.5 Å for full-length human 7,8-DHF-PDXP with phosphate (PDB code 9EM1, colored in wheat yellow, *left panel*) and 1.5 Å for full-length human 7,8-DHF-PDXP without phosphate (PDB code 8S8A, colored in light blue, *right panel*). One protomer of each homodimeric PDXP is shown in cartoon representation and the other protomer in surface representation. 7,8-DHF is displayed in sphere representation with its C-atoms in cyan. Mg^2+^ ions are shown as deep purple spheres and phosphate ions are shown in sphere representation with the phosphorous atom in orange. (**b**) Orientation of 7,8-DHF in the active sites of human 7,8-DHF-PDXP in the presence or absence of phosphate. Structural details of bound 7,8-DHF and adjacent residues of the active sites are shown. *Left,* phosphate-containing 7,8-DHF-PDXP (wheat yellow, cartoon representation). *Right,* phosphate-free 7,8-DHF-PDXP (light blue, cartoon representation). 7,8-DHF is shown in stick representation (cyan C-atoms). The corresponding amino acids in murine PDXP are given in parentheses (see also **Fig. 3 – figure supplement 1e, f**). (**c**) Comparison of the Mg^2+^ coordination spheres. *From left to right*: human apo-PDXP (PDB: 2P27), human PDXP in complex with PLP (PDB: 2CFT), human PDXP in complex with 7,8-DHF in the presence of phosphate (PDB: 9EM1), human PDXP in complex with 7,8-DHF in the absence of phosphate (PDB: 8A8S). The catalytically essential Mg^2+^ is shown as a deep purple sphere. In 2CFT, Mg^2+^ was exchanged for Ca^2+^, which is shown here as a light brown-colored sphere. Water molecules are shown as blue spheres. (**d**) Verification of 7,8-DHF - PDXP interactions. *Left panel*, phosphatase activity of purified PDXP or the indicated PDXP variants. Data are mean values ± S.D. of *n*=3 independent experiments. *Right panel*, determination of the IC_50_ values of 7,8-DHF for purified PDXP or the indicated PDXP variants. Data are mean values ± S.D. of *n*=3 independent experiments. Apparently missing error bars are hidden by the symbols. **Source data** are available for this Figure.

**Table 2.**
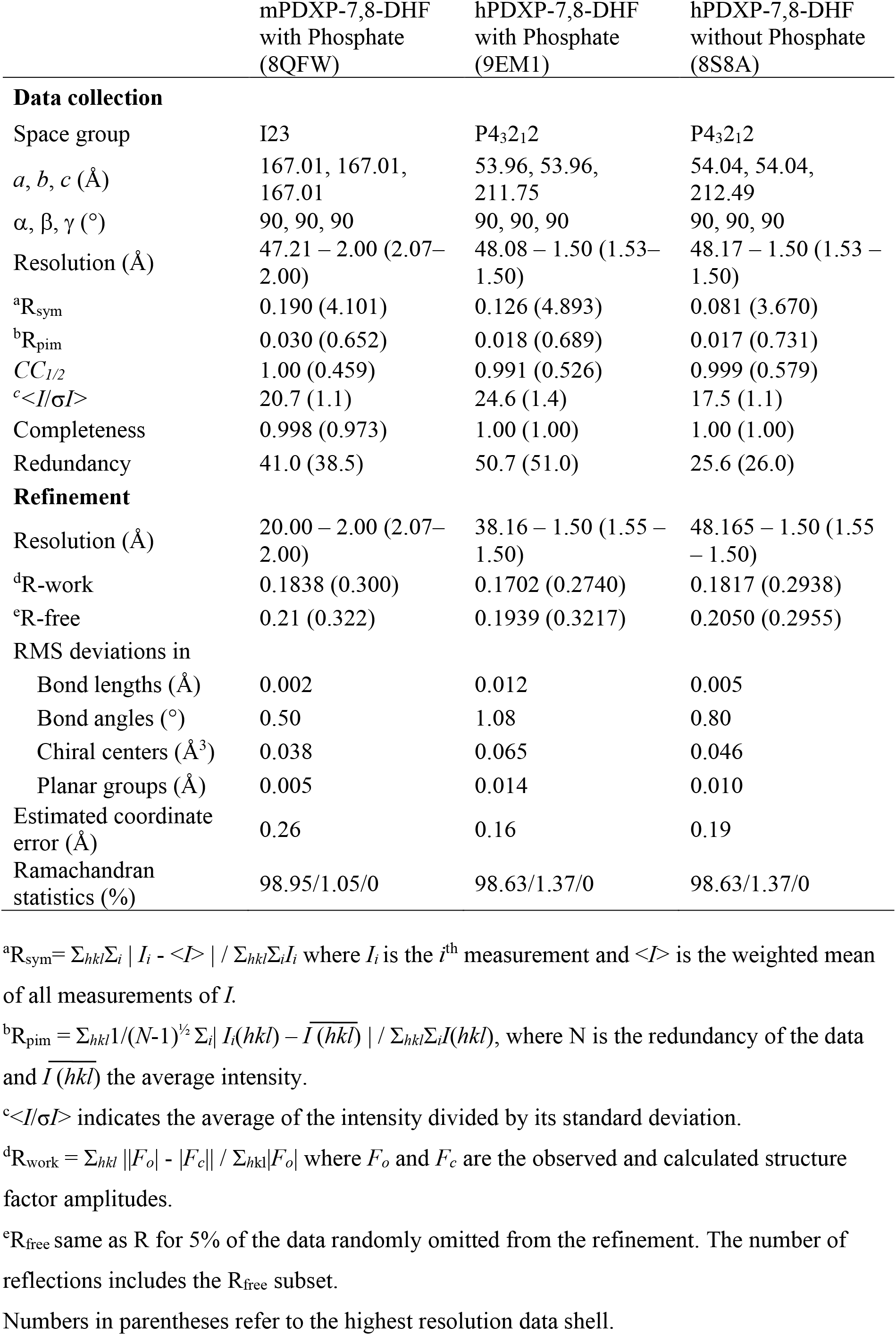
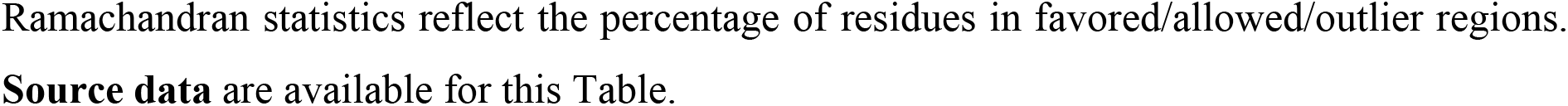
Data Collection and Refinement Statistics.

Like their respective apo-forms, 7,8-DHF-bound murine and human PDXP homodimerize via their cap domains (**Fig. 3a** and **Fig. 3 – figure supplement 1a**). The Cα atom-based alignment of the structures representing murine apo-PDXP and murine 7,8-DHF-bound PDXP resulted in root mean square (RMS) deviations in the range of 0.43-0.71 Å. Even smaller values were obtained when human apo-PDXP, human PLP-bound PDXP and human 7,8-DHF-bound PDXP were superimposed with RMS deviations in the range from 0.29-0.54 Å (**Table 3**). Hence, binding of the inhibitor did not result in significant changes in murine or human PDXP backbone conformations. All catalytic core residues and the Mg^2+^ cofactor are correctly oriented in the presence of the inhibitor. We conclude that 7,8-DHF binding does not appear to impact the overall fold of murine or human PDXP.

**Table 3.** Alignment of murine and human PDXP structures.

|  | 7,8-DHF-mPDXP (+ P),<br>protomer A | 7,8-DHF-mPDXP (- P),<br>protomer B |
| --- | --- | --- |
| 7,8-DHF-mPDXP (+ P), protomer B | 0.43 Å |  |
| Apo-mPDXP, protomer A | 0.50 Å | 0.71 Å |
| Apo-mPDXP, protomer B | 0.60 Å | 0.69 Å |

|  | 7,8-DHF-hPDXP (+ P) | 7,8-DHF-hPDXP (- P) |
| --- | --- | --- |
| 7,8-DHF-hPDXP (- P) | 0.37 Å |  |
| Apo-hPDXP | 0.54 Å | 0.45 Å |
| PLP-hPDXP | 0.39 Å | 0.29 Å |

Cα atom-based alignment of the structures representing murine apo-PDXP (PDB: 4BX3), 7,8-DHF-bound murine PDXP (with inhibitor-bound protomer A and inhibitor-free protomer B; PDB: 8QFW), human apo-PDXP (PDB: 2P27), 7,8-DHF-bound human PDXP with phosphate (+ P) (PDB: 9EM1), 7,8-DHF-bound human PDXP without phosphate (- P) (PDB: 8S8A) and PLP-bound human PDXP (PDB: 2CFT); mPDXP, murine PDXP; hPDXP, human PDXP. Root mean square deviations are indicated.

7,8-DHF was observed to only bind to one subunit (the A-chain) of murine PDXP (**Fig. 3 – figure supplement 1a**) with well-defined density (**Fig. 3 – figure supplement 1b**) and full occupancy since its average B-factor of 45.8 Å^2^ closely matches the B-factors of the surrounding atoms. Binding to the other subunit (B-protomer) is prevented by a salt bridge between Arg62 and Asp14 of a symmetry-related A-protomer in the crystal (**Fig. 3 – figure supplement 1c**). The χ_1_ and χ_2_ torsion angles of the Arg62 side chain observed in the B-protomer correspond to those observed for this side chain in both protomers of the murine apo-structure (4BX3). To allow binding of the inhibitor, the side chain of Arg62 needs to adopt a completely extended conformation, which is prevented by the salt bridge. However, preventing mPDXP salt bridge formation by mutating Asp14 to Ala did not alter the efficacy of 7,8-DHF inhibition (**Fig. 3 – figure supplement 1d**; see also **Fig. 3d** for the characterization of the PDXP-Arg62Ala variant). It is therefore currently unclear whether the mPDXP crystal state with only a single inhibitor bound per dimer reflects the state in solution. Due to the limited solubility of 7,8-DHF, we were unable to address the stoichiometry of 7,8-DHF binding to the PDXP dimer with isothermal calorimetry. It is conceivable that the mPDXP crystal packing is very stable (indeed, 7,8-DHF-bound mPDXP crystallized in the same cubic space group as apo-mPDXP, see ref. (48), including the aforementioned salt bridge between Arg62 of the B-subunit and Asp14 of a symmetry-related molecule), and that the free energy generated by the formation of the crystal lattice is higher than the free energy generated upon inhibitor binding.

In contrast to murine PDXP, 7,8-DHF bound to human PDXP with a ratio of two inhibitors per homodimer (**Fig. 3a**) and well-defined density (**Fig. 3 – figure supplement 2a**). Interestingly, the orientation of the inhibitor was markedly affected by the presence or absence of phosphate (**Fig. 3b**). In the presence of phosphate, the inhibitor moiety that is closest to the Mg^2+^ cofactor is the uncharged phenyl ring of 7,8-DHF. In the absence of phosphate, the inhibitor is flipped horizontally, with the hydroxylated chromone substructure of 7,8-DHF now located closest to the Mg^2+^ ion (**Fig. 3b**, compare *left* and *right panels*). The inhibitor localization in the presence of phosphate was identical in human and murine PDXP (**Fig. 3 – figure supplement 2e**). The localization of the phosphate ion that co-crystallized with 7,8-DHF-bound human or murine PDXP overlaps exactly with the localization of the PLP phosphate moiety introduced from PDB code 2CFT (human PDXP in complex with PLP) for visualization purposes (**Fig. 3c** and **Fig. 3 – figure supplement 2b**), indicating that the phosphate ion is bound in a catalytically relevant position.

Irrespective of the orientation of 7,8-DHF in the PDXP active site, the inhibitor is embedded in a cavity that is exclusively formed by the active site of protomer A, without a contribution of the dimerization interface with protomer B. All PDXP residues found to engage in 7,8-DHF interactions are identical in murine and human PDXP (**Fig. 3 – figure supplement 3**). One side of this cavity is formed by the more polar residues Asp27, Asn60, Ser61 and Arg62 (identical amino acid residue numbering in mPDXP and hPDXP), whereas the opposite side is established by the more hydrophobic residues Tyr150, His182, Pro183 and Leu184 (corresponding to Tyr146, His178, Pro179, and Leu180 in mPDXP). Adjacent to this hydrophobic stretch, the polar residue Glu152 (Glu148 in mPDXP) is located at the active site entrance, directly opposite of Arg62 on the more polar side of the 7,8-DHF binding channel (**Fig. 3b** and **Fig. 3 – figure supplement 1e**).

Interestingly, Glu152 (or Glu148) and Arg62 can form an intramolecular salt bridge that obstructs the active site entrance (**Fig. 3 – figure supplement 4**). This interaction was observed in phosphate-free 7,8-DHF-hPDXP and phosphate-free PLP-hPDXP, as well as in apo-hPDXP and apo-mPDXP. In contrast, the 7,8-DHF binding pose that is dictated by the concomitant binding of phosphate and 7,8-DHF interferes with the Glu152 (Glu148)-Arg62 interaction in both, hPDXP and mPDXP (**Fig. 3 – figure supplement 4**). Thus, although we did not find evidence for major cap/core or substrate-specificity loop movements (40, 48) in PDXP, the presence or absence of a salt bridge formed between the cap domain residue Glu152 (Glu148) and the core domain residue Arg62 indicates subtle conformational changes in PDXP that may mediate an opening or a closure of the active site entrance.

Inhibitor binding in the presence of phosphate is identical in human and murine PDXP (**Fig. 3 – figure supplement 1f**) and appears to be primarily stabilized by two hydrogen bonds, as well as polar and non-polar interactions (**Fig. 3b**, *left panel*). The side chain hydroxyl group of Ser61 forms a direct hydrogen bond with the ketone group of the inhibitor, which is additionally coordinated by the Ser61 backbone nitrogen atom. Furthermore, Glu152 (Glu148) forms a direct hydrogen bond via its carboxylic acid with the 7-hydroxyl group of 7,8-DHF. The side chains of the polar residues Asp27, Asn60 and Arg62 engage in van der Waals interactions with 7,8-DHF. The two hydroxyl groups of the 7,8-DHF benzyl ring engage in van der Waals interactions with the guanidinium group of Arg62 and the carboxylic acid function of Glu148 (Glu152). On the more hydrophobic side of the binding cavity, Tyr146 (Tyr150) forms π-electron stacking interactions with the pyrone ring of 7,8-DHF. In addition, the His178 (His182) imidazole group coordinates the 7,8-DHF phenyl ring via a cation-π interaction. His178 (His182), located in the substrate specificity loop, and Asn60 and Arg62 are also important for PLP binding (48, 49).

Inhibitor binding in the absence of phosphate is primarily stabilized by metal coordination and hydrogen bonds, as well as polar and non-polar interactions (**Fig. 3b**, *right panel*). The 7-hydroxyl group of 7,8-DHF is involved in an octahedral Mg^2+^ coordination, albeit with an elongated oxygen-Mg^2+^ distance of 2.7 Å, leading to the displacement of a water molecule (**Fig. 3c**). This inhibitor-based water displacement is not observed in the phosphate-containing murine or human 7,8-DHF-PDXP structures. The 8-hydroxyl group of 7,8-DHF forms a hydrogen bond with Asp239 and His182. The ketone group of the inhibitor participates in a water-bridged hydrogen bond to the carboxyl group of Asp27 and the backbone amine group of Gly33. Like in the phosphate-containing structure, the His182 imidazole group coordinates the 7,8-DHF phenyl ring via π-π stacking. In addition, Tyr150 forms edge to face π-π stacking interactions with the phenyl ring of 7,8-DHF. The side chains of the polar residues Asp27, Asn60, and, to some degree, also of Arg62, engage in van der Waals interactions with 7,8-DHF.

To verify the putative 7,8-DHF – PDXP interactions, we introduced single mutations into the binding interface. Asn60, Arg62, Tyr146, Glu148 and His178 in mPDXP were each exchanged for Ala (PDXP^N60A^, PDXP^R62A^, PDXP^Y146A^, PDXP^E148A^, or PDXP^H178A^, respectively). Since the carboxamide group of Asn60 can form a hydrogen bond with the carboxylate moiety of Asp27, and a loss of this interaction in the PDXP^N60A^ variant is predicted to alter the PDXP structure, we additionally mutated Asn60 to Ser (PDXP^N60S^). PDXP variants were recombinantly expressed and purified from *E. coli* (see **Fig. 3 – figure supplement 5** for protein purity). **Figure 3d** (left panel) shows that all PDXP variants were enzymatically active. As expected, the phosphatase activities of PDXP^N60A^ and of PDXP^H178A^ were reduced. The somewhat elevated phosphatase activity of PDXP^Y146A^ and PDXP^E148A^ is currently unexplained. Importantly, all variants except PDXP^N60A^ and PDXP^N60S^ were resistant to 7,8-DHF, supporting the essential role of each of these residues for inhibitor binding and the minor contribution of the weak van der Waals interactions between Asn60 and 7,8-DHF during inhibitor binding (**Fig. 3d**, right panel). These data also suggest that Asn61 contributes to the limited efficacy of 7,8-mediated PDXP inhibition in vitro.

Based on the inhibitor-bound structures and the predominantly competitive component of PDXP inhibition by 7,8-DHF (increased K_M_, see **Table 1**), it seems likely that 7,8-DHF sterically hinders substrate access to the active site, and competes with PLP coordination (**Fig. 3 – figure supplement 2b**). In addition, BLI measurements (see **Fig. 2c**) showed a relatively slow association rate and extended residence time of 7,8-DHF (τ=30.3 s). This may indicate a reorganization of the Mg^2+^-coordination due to inhibitor binding, and a reorientation of 7,8-DHF during the PDXP catalytic cycle. The reduced rate of product formation may account for the apparent mixed mode of 7,8-DHF-mediated PDXP inhibition (reduction of v_max_, see **Table 1**).

### 7,8-DHF functions as a PDXP inhibitor in hippocampal neurons

To investigate cellular target engagement of 7,8-DHF, we isolated primary hippocampal neurons from PDXP-WT and PDXP-KO embryos. PDXP deficiency increased total PLP levels 2.4-fold compared to PDXP-WT neurons (**Fig. 4a**). This finding is in good agreement with the PLP increase resulting from PDXP loss in total hippocampal extracts (see **Fig. 1**). The larger absolute PLP values in cultured neurons are likely attributable to the high concentration of the PLP precursor pyridoxal (20 µM) in the culture medium. We did not observe PDXP-dependent changes in pyridoxal kinase (PDXK) expression (**Fig. 4b**) and could not detect pyridox(am)ine-5’-phosphate oxidase (PNPO) in hippocampal neuronal cultures, suggesting that the PLP increase was primarily caused by the constitutive PDXP loss.

**Figure 4.**
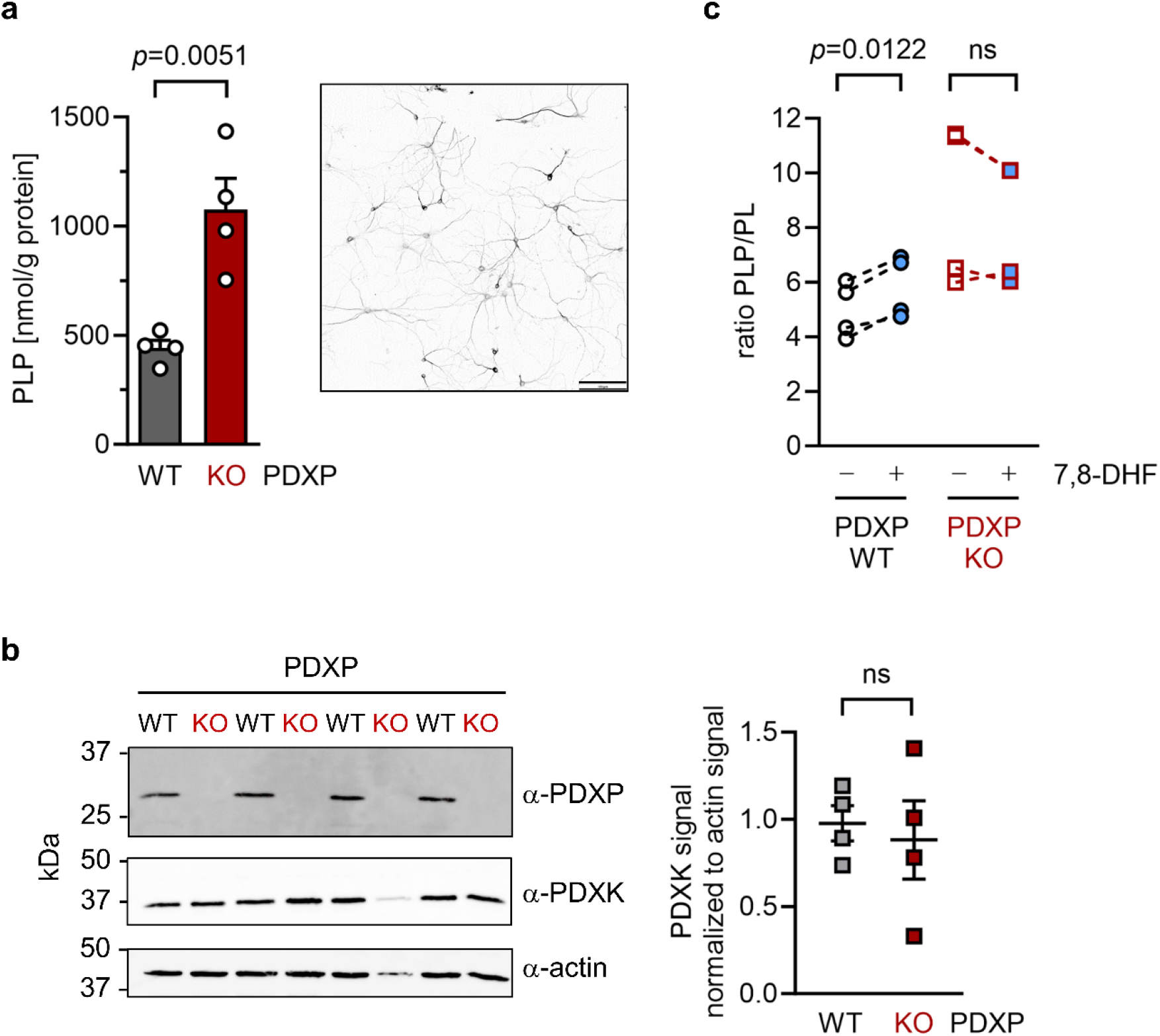
Effect of 7,8-DHF on the PLP/PL ratio in cultured hippocampal neurons from WT or PDXP-KO mice. (**a**) Effect of long-term PDXP deficiency on total PLP levels in hippocampal neurons. Data are mean values ± S.E. of *n*=4 independent experiments. Statistical significance was assessed with a two-tailed, unpaired *t*-test. A representative image of primary hippocampal neurons stained for the neuronal marker protein MAP2 is shown in the insert (pixel intensities were color-inverted for better visualization). Scale bar, 100 µm. (**b**) Western blot analysis of PDXP and PDXK expression in hippocampal neuron samples shown in (a). The same blots were reprobed with α-actin antibodies as a loading control. The densitometric quantification of PDXK signals is shown on the right; data are mean values ± S.E. of *n*=4 independent experiments. (**c**) Effect of 7,8-DHF (20 µM, 45 min) or the DMSO solvent control (0.02% v/v, 45 min) on the PLP/PL ratio in hippocampal neurons of PDXP-WT or PDXP-KO mice. **Source data** are available for this Figure.

To assess the consequences of 7,8-DHF treatment on PLP levels in hippocampal neurons, we chose short-term incubation conditions (45 min, 20 µM) to avoid possible secondary effects of the inhibitor. As expected, the acute effect of 7,8-DHF treatment in WT cells was much more subtle (⁓9% increase in total PLP) than the impact of long-term PDXP-deficiency (441.3 ± 62.6 nmol PLP/g protein in DMSO solvent control-treated cells versus 482.7 ± 130.4 nmol PLP/g protein in 7,8-DHF treated cells; data are mean values ± S.E. of *n*=4 independent experiments). However, this effect is likely underestimated because only the PDXP-accessible pool of non-protein-bound PLP may be impacted by 7,8-DHF (see **Fig. 1c**). Due to the limited number of available hippocampal neurons, we were unfortunately unable to obtain sufficient quantities of protein-depleted PLP pools to address this question.

Acute changes in the PLP/PL ratio may be a more sensitive indicator of PDXP activity than changes in total PLP levels alone, because PDXP inhibition is expected to increase cellular levels of PLP (the PDXP substrate) and to concomitantly decrease the levels of PL (the product of PDXP phosphatase activity). The PLP/PL ratio is also independent of the exact protein concentration in a given extract of hippocampal neurons, thus optimizing comparability between samples. As shown in **Fig. 4c**, 7,8-DHF significantly increased the PLP/PL ratio in PDXP-WT, but not in PDXP-KO hippocampal neurons (+18% versus +1% compared to the respective DMSO controls). Together, these data indicate that 7,8-DHF can modulate cellular PLP levels in a PDXP-dependent manner and validate PDXP as a 7,8-DHF target in primary hippocampal neurons.

## DISCUSSION

PLP deficiency has been associated with human brain disorders for decades (3), yet causal links remain unclear. Aside from vitamin B6 administration, pharmacological strategies to elevate intracellular PLP levels are lacking. Here, we identify 7,8-dihydroxyflavone (7,8-DHF) as a direct PDXP inhibitor that increases PLP levels in hippocampal neurons, validating PDXP as a druggable target to control intracellular PLP levels in the brain. We also present three high-resolution 7,8-DHF/PDXP co-crystal structures that will facilitate the design of more potent, efficacious, and selective PDXP inhibitors in the future. Such molecules might improve the control of intracellular PLP levels and help to elucidate a possible contribution of PLP to the pathophysiology of brain disorders. Our observation that the expression of PDXP is substantially upregulated in hippocampi of middle-aged mice suggests that a therapeutic vitamin B6 supplementation alone may not suffice to elevate intracellular PLP levels under conditions where the PLP-degrading phosphatase is hyperactive.

The discovery of 7,8-DHF as a direct PDXP inhibitor was unexpected. Interestingly, numerous in vivo studies have reported the effectiveness of 7,8-DHF in brain disorder models, including rodent models of Alzheimer’s disease (50–57), depression (58–63), schizophrenia (64–68), epilepsy (69, 70) and autism (71–74). Although PLP deficiency is thought to contribute to the respective human conditions (3, 75, 76), PLP-dependent processes have not yet been considered in the context of 7,8-DHF-induced effects.

7,8-DHF was initially discovered as a small-molecule TrkB agonist with BDNF-mimetic activity (77). BDNF, a high-affinity TrkB ligand, is an important neuropeptide for nervous system function and pathology. Consensus is emerging that BDNF plays a key role in the treatment response to neuropsychiatric drugs (32). Therapeutics that target BDNF/TrkB-signaling are thus of interest as disease-modifying agents in several brain disorders. Since BDNF does not cross the blood-brain barrier, attempts have been made to develop small molecule BDNF mimetics. Several candidates have been reported, including 7,8-DHF (33, 77). Nevertheless, the on-target selectivity and efficacy of these compounds is actively debated. Using quantitative and direct assays to measure TrkB dimerization and activation, TrkB downstream signaling pathways, TrkB-dependent gene expression and cytoprotection, 7,8-DHF and other reported small-molecule TrkB agonists failed to activate TrkB in cells (33–35). An electrophysiological study in acute hippocampal slice preparations demonstrated that 7,8-DHF potentiates hippocampal mossy fiber-CA3 synaptic transmission in a TrkB receptor-independent manner (78). Overall, it appears that the mechanism of action of 7,8-DHF is incompletely understood, but 7,8-DHF targets other than TrkB so far have remained elusive. The identification of 7,8-DHF as a PDXP inhibitor reported here indicates that this flavone may modulate vitamin B6-dependent processes and suggests that PDXP could be explored as a pharmacological entry point into brain disorders.

## MATERIALS AND METHODS

### Materials

Unless otherwise specified, all reagents were of the highest available purity and purchased from Sigma Aldrich (Schnelldorf, Germany). 3,7,8,4’-Tetrahydroxyflavone was obtained from Ambinter (Orléans, France), all other flavones were from Sigma Aldrich.

### PDXP knockout mice

Floxed PDXP mice (*Pdxp^tm1Goh^*) were generated on a C57Bl/6J background, and whole-body *Pdxp* knockouts were achieved by breeding with B6.FVB-Tg(EIIa-cre)C5379Lmgd/J (EIIa-Cre) transgenic mice, as described (30). All experiments were authorized by the local veterinary authority and committee on the ethics of animal experiments (Regierung von Unterfranken). All analyses were carried out in strict accordance with all German and European Union applicable laws and regulations concerning care and use of laboratory animals.

### Preparation of hippocampi and hippocampal neurons and immunocytochemistry

Mice were sacrificed by cervical dislocation, and brains were immediately placed on a pre-cooled metal plate and dissected under a Leica M80 binocular (Leica, Wetzlar, Germany). Hippocampi were weighed and flash-frozen in liquid nitrogen. The entire procedure was performed in <3 min. Hippocampal lysates were prepared by the addition of ice-cold PBS (200 µL PBS/10 mg hippocampal wet weight) and homogenized for 1 min in a TissueLyser II instrument (Qiagen, Hilden, Germany). One fourth of the obtained volume of each lysate was used for the analysis of total PLP concentrations as described below. To determine protein-depleted PLP (27), the remaining volume of each lysate was centrifuged at 14,000 × *g* for 15 min at 4°C. The supernatant was applied to 3 kDa MWCO filters (Amicon Ultra-0.5 Centrifugal Filter; Merck Millipore, Darmstadt, Germany), and centrifuged at 14,000 × *g* for 45 min at 4°C. The flow-through was collected and prepared for HPLC analysis (see below).

Primary hippocampal neuronal cultures were prepared from mouse embryos at embryonic day 17. Hippocampi were incubated with 0.5 mg/mL trypsin, 0.2 mg/mL EDTA and 10 µg/mL DNase I in PBS for 30 min at 37°C. Trypsinization was stopped by adding 10% fetal calf serum. Cells were dissociated by trituration, counted, and seeded at a density of 150,000 cells per 35 mm dish. Dissociated cells were grown in neurobasal medium supplemented with L-glutamine and B27 supplement (A3582801, Life Technologies, Dreieich, Germany) with an exchange of 50% of the medium after 6 days in culture. After 21 days of differentiation (day in vitro 21/DIV21), 7,8-DHF (20 µM) or DMSO (0.02%, v/v) was added to the hippocampal neuronal cultures for 45 min. Cells were rinsed once with PBS (37°C), lysed in 150 µL ice cold H_2_O, and placed at −80°C for at least 30 min.

For immunocytochemistry, DIV21 primary hippocampal neurons were fixed with 4% (w/v) paraformaldehyde in phosphate-buffered saline (PBS) for 15 min at RT. After washing twice with PBS, 50 mM NH_4_Cl was added for 10 min. Cells were then permeabilized with 0.1% (v/v) Triton X-100 and blocked with 5% (v/v) goat serum in PBS for 30 min at 22°C. Cells were incubated with mouse monoclonal anti-MAP2 antibodies (1:500 dilution, clone MAB3418, Millipore, Darmstadt, Germany) for 1 h in 5% goat serum/PBS at 22°C. Alexa-488-labeled secondary goat anti-mouse antibodies (1:500 dilution; Dianova, Hamburg, Germany) were applied for 1 h. Nuclei were counter-stained with 4’,6-diamino-2-phenylindole (DAPI), and slides were mounted with Mowiol. Images were acquired using an inverted IX81 microscope equipped with an Olympus UPLSAPO 60× oil objective (numerical aperture: 1.35) on an Olympus FV1000 confocal laser scanning system, using a FVD10 SPD spectral detector and diode lasers of 405 nm (DAPI) and 495 nm (Alexa488).

### Determination of PLP and PL by high-performance liquid chromatography (HPLC)

Samples were derivatized as described (79). Briefly, 100 µL of lysate were mixed with 8 µL derivatization agent (containing 250 mg/mL of both semicarbazide and glycine), and incubated on ice for 30 min. Samples were then deproteinized by addition of perchloric acid (8 µL of a 72% (w/v) stock solution), followed by centrifugation at 15,000 × *g* for 15 min at 4 °C. Supernatants (100 µL) were neutralized with 10 µL NaOH (25% (v/v) stock solution), and 2 µM pyridoxic acid was added as an internal standard. PLP and PL were subjected to the same derivatization protocol to establish a standard curve. Samples were analyzed on a Dionex Ultimate 3000 HPLC (Thermo Fisher Scientific, Dreieich, Germany), using 60 mM Na_2_HPO_4_, 1 mM EDTA, 9.5% (v/v) MeOH; pH 6.5 as mobile phase. PL, PLP and pyridoxic acid were separated on a 3 μm reverse phase column (BDS-HYPERSIL-C18, Thermo Fisher Scientific). Chromatograms were analyzed using Chromeleon 7 software (Thermo Fisher Scientific).

### Western blotting

Tissue or cell homogenates (prepared as detailed above for HPLC analysis) were extracted with 4 × RIPA buffer (final concentration, 50 mM Tris, pH 7.5; 150 mM NaCl, 1% (v/v) Triton X-100, 0.5% (v/v) sodium deoxycholate, 0.1% (w/v) SDS, 1 mM 4-(2-aminoethyl)benzenesulfonyl fluoride (Pefabloc), 5 μg/mL aprotinin, 1 μg/mL leupeptin, 1 μg/mL pepstatin) for 15 min at 4 °C under rotation, and lysates were clarified by centrifugation (20,000 × *g*, 15 min, 4 °C). Protein concentrations in the supernatants were determined using the Micro BCA Protein Assay Kit (Thermo Fisher Scientific). Proteins were separated by SDS-PAGE and transferred to nitrocellulose membranes by semidry-blotting. Antibodies were purchased from the following providers: Merck Millipore (mouse monoclonal α-actin mAb1501, dilution 1:5000); Cell Signaling (rabbit monoclonal α-PDXP clone C85E3, dilution 1:1000; Cell Signaling, Danvers/Massachusetts, USA), Sigma Aldrich (rabbit polyclonal α-PDXK/AB1, #AV53615, dilution 1:1000), and Thermo Fisher Scientific (rabbit polyclonal α-PNPO, #PA5-26400, dilution 1:1000, as used in ref. (30)). Western blots were densitometrically quantified with NIH ImageJ, version 1.45i.

### Phosphatase plasmids and cloning

N-terminally GST-tagged, human PDXP was in pGEX-4T-1 (Amersham Biosciences, Amersham, UK). N-terminally His_6_-SUMO-tagged human PDXP was cloned into pET-SUMO (coding for human SUMO; a kind gift of Dr. Pedro Friedmann Angeli, Rudolf-Virchow-Center, University of Würzburg, Germany); His_6_-SenP2 (EMBL Heidelberg) was in pET-M11. All other phosphatases were of murine origin and were subcloned into pET-M11 (EMBL), as described (40). *Pdxp* point mutants (generated by nested PCR) were subcloned into the *Nco*I (*Pci*I for *Psph*) and *EcoR*I restriction sites of pET-M11, using Q5 Hot Start High-Fidelity DNA Polymerase (New England Biolabs, Frankfurt/Main, Germany). *Pdxp-D14N* was generated with the Platinum SuperFi II DNA Polymerase Mastermix according to mutagenesis protocol A provided by the manufacturer (Thermo Fisher Scientific) and cloned into pET-M11 as described above. The following primers (oligonucleotide sequence 5’-3’; fwd, forward; rev, reverse) were used:

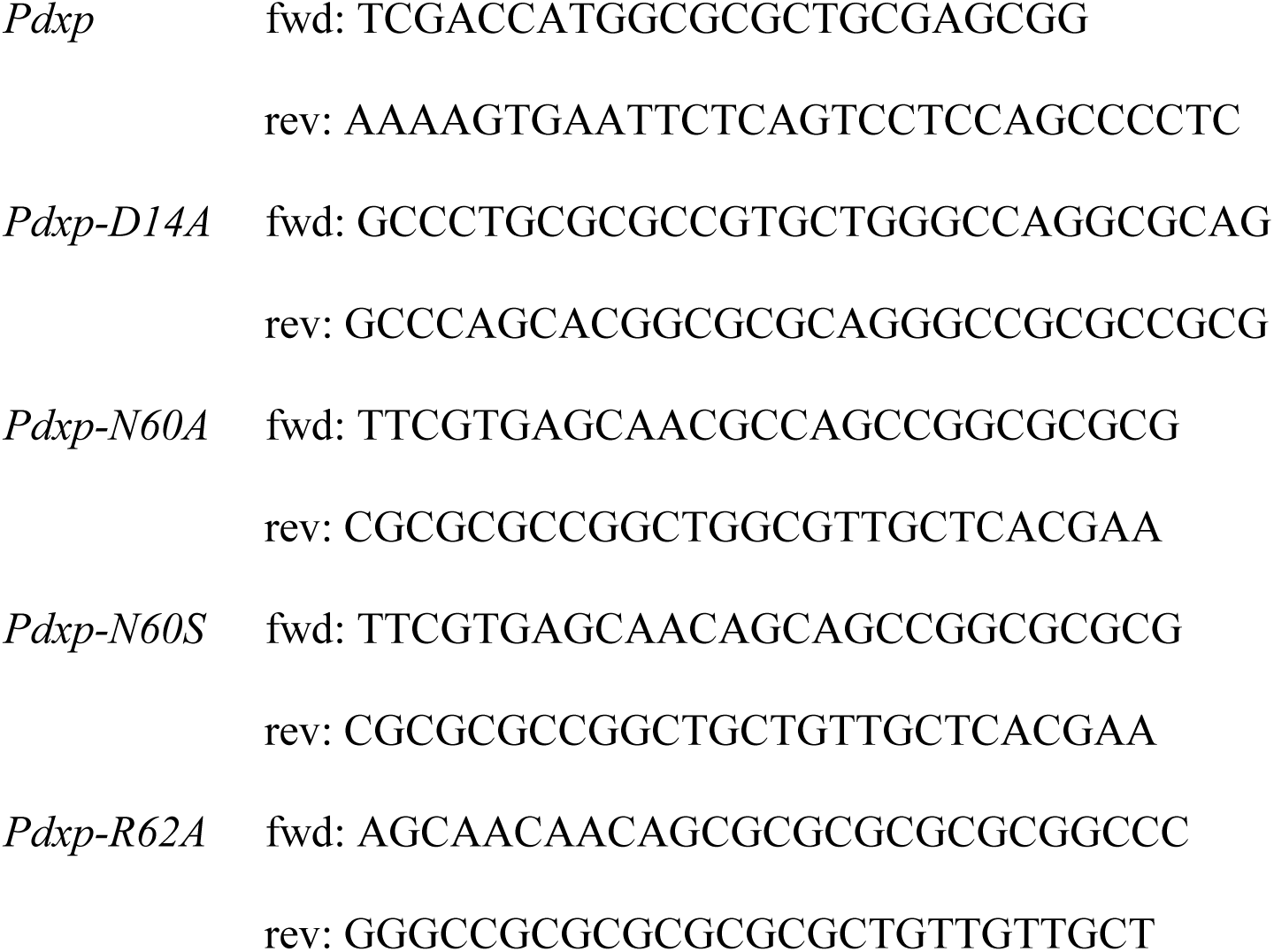
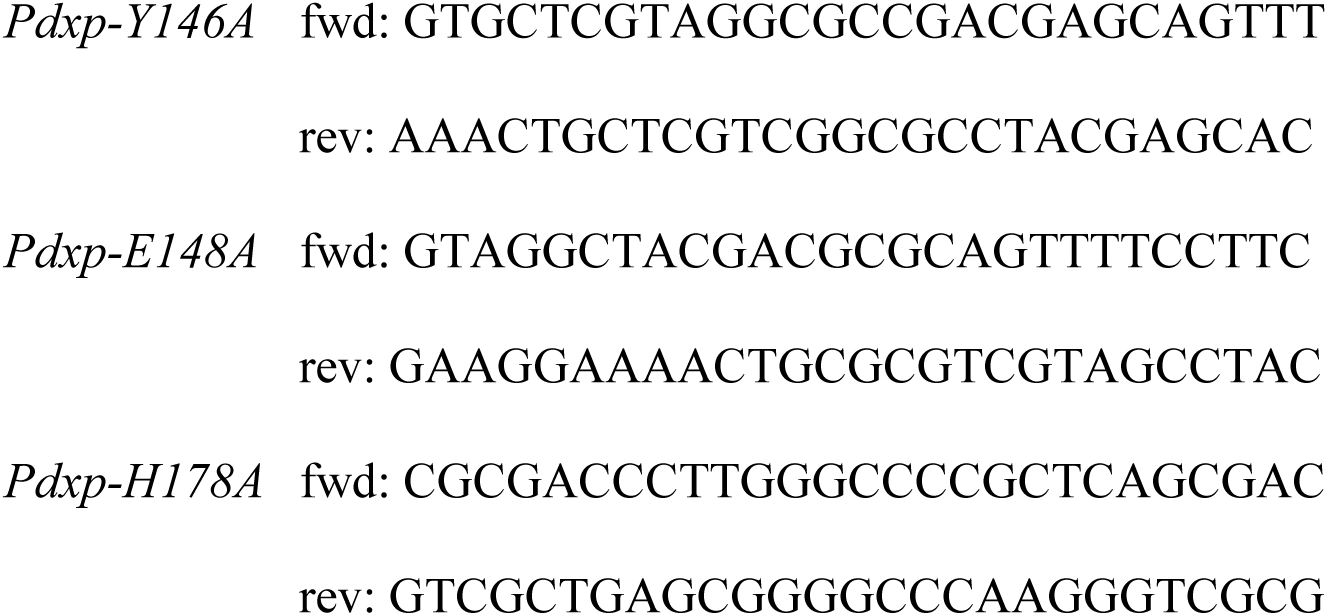

Primers were purchased from Eurofins Genomics (Ebersberg, Germany), and all constructs were verified by sequencing (Microsynth Seqlab, Göttingen, Germany).

### Expression and purification of recombinant proteins

Human His_6_-SUMO-tagged PDXP was grown in ZYP-5052 autoinduction medium for 7 h at 37° C, followed by 48 h at 21°C (80). All purification steps of murine PDXP and murine PGP were carried out exactly as described (40). The purification of the His_6_-SUMO-tagged human PDXP was carried out exactly as described for murine His_6_-tagged PDXP (40), except that human SenP2 protease was used to cleave the His_6_-SUMO-tag. N-terminally His_6_-tagged PDXP variants and His_6_-SenP2 were expressed as described for PDXP-WT (40).With the exception of PDXP-D14, the His_6_-tag was not cleaved off. Human GST-PDXP was transformed into *E. coli* BL21(DE3) (Stratagene Europe/VWR, Darmstadt, Germany). Protein expression was induced with 0.5 mM isopropyl β-d-thiogalactopyranoside for 18 h at 20 °C. All subsequent purification steps were carried out at 4 °C. Cells were harvested by centrifugation for 10 min at 8000 × *g* and resuspended in lysis buffer (100 mM triethanolamine/TEA, 500 mM NaCl; pH 7.4) supplemented with protease inhibitors (EDTA-free protease inhibitor tablets; Roche, Mannheim, Germany) and 150 U/mL DNase I (Applichem, Renningen, Germany). Cells were lysed using a cell disruptor (Constant Systems, Daventry, UK), and cell debris was removed by centrifugation for 30 min at 30,000 × *g*. GST-PDXP was batch-purified on a glutathione sepharose 4B resin (GE Healthcare, Uppsala, Sweden). After extensive washing with 25 column volumes of wash buffer (50 mM TEA, 250 mM NaCl; pH 7.4), GST-PDXP was eluted in wash buffer supplemented with 10 mM reduced glutathione, concentrated, and further purified in buffer A (50 mM TEA, 250 mM NaCl, 5 mM MgCl_2_; pH 7.4) using a HiLoad 16/60 Superdex 200 pg gel filtration column operated on an ÄKTA liquid chromatography system (GE Healthcare).

### High-Throughput Screen for PDXP Modulators

The screening campaign (chemical library, screening protocol, concentration-dependent assays, data analysis) was conducted exactly as described previously (40), except that the primary screen was done with PDXP, the counter-screen with PGP, and PDXP inhibitor hits were validated using 5’-pyridoxal phosphate (PLP) as a physiological PDXP substrate.

### IC_50_ determinations, enzyme kinetics, and compound selectivity

Conditions for enzymatic assays were as previously published (40), with the following modifications. Bovine brain calcineurin (PP2B, Sigma Aldrich #C1907) activity against the PKA regulatory subunit type II (phosphopeptide DLDVPIPGRFDRRVpSVAAE; Sigma Aldrich #207008) was assayed at 37°C in 100 mM NaCl, 50 mM Tris, 6 mM MgCl_2_, 0.5 mM CaCl_2_, 0.5 mM DTT, 0.025% (v/v) NP40; pH 7.5. Recombinant human PTP1B (amino acids 1-321, Cayman Chemical, Ann Arbor, Michigan, USA) activity against the Tyr^992^ autophosphorylation site of EGFR (DADEpYLIPQQG; Santa Cruz Biotechnology, Heidelberg, Germany) was assayed at 30°C in 150 mM NaCl, 50 mM 2-(N-morpholino)ethanesulfonic acid, 1 mM EDTA; pH 7.2. Murine PDXP-D14A was assayed exactly like PDXP-WT in 30 mM triethanolamine, 5 mM MgCl_2_, 30 mM NaCl; pH 7.5, supplemented with 0.01% (v/v) Triton X-100.

Flavone stocks were prepared at 10 mM in 100% DMSO. A constant final DMSO concentration of 0.4% was maintained under all conditions, and solvent control samples contained 0.4% DMSO without compounds. Purified phosphatases were pre-incubated for 10 min at RT with serial dilutions of flavones. Dephosphorylation reactions were started by the addition of the indicated substrate; buffer with substrate and the respective flavone but without the enzyme served as a background control. Prior to compound testing, time courses of inorganic phosphate release from the respective phosphatase substrates were conducted to ensure assay linearity. Inorganic phosphate release was detected with a malachite green solution (Biomol Green; Enzo Life Sciences, Lörrach, Germany); the absorbance at 620 nm (A_620_) was measured on an Envision 2104 multilabel reader (Perkin Elmer, Rodgau, Germany). Released phosphate was determined by converting the values to nmol P_i_ with a phosphate standard curve. Data were analyzed with GraphPad Prism version 9.5.1 (GraphPad, Boston/Massachusetts, USA). For IC_50_ determinations, log_inhibitor_ versus response was calculated (four parameter). To derive K_M_ and k_cat_ values, data were fitted by nonlinear regression to the Michaelis-Menten equation.

### Biolayer interferometry (BLI)

PDXP was biotinylated using the EZ-Link NHS-PEG4-Biotin kit, as recommended by the manufacturer (Thermo Fisher Scientific), and loaded on Super Streptavidin Biosensors (SSA) (Sartorius, Göttingen, Germany) as follows. SSA sensors were equilibrated for 1 h at RT in BLI assay buffer (250 mM triethanolamine, 5 mM MgCl_2_, 250 mM NaCl, 0.005% (v/v) TWEEN-20; pH 7.5), loaded with 200 µg/mL biotinylated PDXP, blocked with 2 µg/mL biocytin, and washed in BLI assay buffer. Reference SSA sensors were blocked with 2 µg/mL biocytin (81). Six point 1:1 serial dilution series of 7,8-DHF and 5,7-DHF were prepared in DMSO, and BLI assay buffer was added to the wells to obtain a 7,8-DHF starting concentration of 25 µM. The final DMSO concentration was 5% (v/v). Buffers for baseline, dissociation, and buffer correction wells were supplemented with the same amount of DMSO for identical buffer conditions. Four measurements were carried out per condition, using one sensor set for two measurements. All measurements were conducted on an Octet K2 device (Sartorius) using 96-well plates. Assay settings were as follows: baseline measurement 45 sec, association time 90 sec, dissociation time 150 sec. The resulting data were processed using the double reference method of the Octet analysis software for removal of drifts and well-to-well artefacts. Kinetic analyses were performed using the Octet analysis software. The steady state analysis was carried out with OriginPro 2021b (OriginLab, Northampton/Massachusetts, USA), using a dose-response model for regression. Due to the poor solubility of 7,8-DHF, the highest concentration of 25 µM was not included in the analysis.

### PDXP crystallization and data collection

For co-crystallization with 7,8-DHF, full-length murine PDXP (10 mg/mL in 50 mM triethanolamine; 250 mM NaCl; 5 mM MgCl_2_; pH 7.4) was supplemented with a three-fold molar excess of the flavone. Prism-shaped crystals of 7,8-DHF-bound murine PDXP were grown at 20°C in 0.1 M phosphate citrate (pH 4.2) and 40% (v/v) PEG 300 using the sitting-drop vapor diffusion method. Human PDXP crystals were grown at 20 °C in 0.1 M Tris (pH 8.5) and 1 M diammonium hydrogen phosphate, or in 0.1 M HEPES (pH 7.0), 15% (v/v) Tacsimat pH 7.0 (Hampton Research, Aliso Viejo, USA) and 2% (w/v) PEG 3350 using the sitting-drop vapor diffusion method. Crystals were cryoprotected for flash-cooling in liquid nitrogen by soaking in mother liquor containing 25% (v/v) glycerol. Diffraction data of murine PDXP in complex with 7,8-DHF were collected from flash-cooled crystals at a temperature of 100 K on beamline BL 14.1 at the BESSY synchrotron (Helmholtz Zentrum Berlin, Germany). Diffraction data of 7,8-DHF bound to human PDXP were collected on beamline ID23-2 at the ESRF (Grenoble, France) [https://data.esrf.fr/doi/10.15151/ESRF-ES-1409594895]. Diffraction data were processed using XDS (82) and further analysed with Aimless (83) of the CCP4 suite (84). The structures of 7,8-DHF-PDXP were solved by molecular replacement with the program Phaser (85) with the structure of the murine PDXP (PDB entry 4BX3) or human PDXP (PDB entry 2P27) as search models, and refined with Phenix (86). Model building was carried out in COOT (87). Structural illustrations were prepared with PyMOL 2.5.1 (88).

## ACKNOWLEDGMENTS

We thank Carola Seyffarth and Nicole Bader for excellent technical assistance, Dr. Jochen Kuper for collecting the murine PDXP diffraction data, the staff at beamline BL14.1 of the BESSY synchrotron, and the staff at beamline ID23-2 of the ESFR synchrotron for technical support. A part of this work was initially funded by the DFG Collaborative Research Center SFB688 (TP A11 to A.G.).

## DATA AVAILABILITY STATEMENT

The previously published PDB entry 4BX3 of murine apo-PDXP [http://doi.org/10.2210/pdb4BX3/pdb], 2P27 of human apo-PDXP [http://doi.org/10.2210/pdb2P27/pdb] and 2CFT of PLP-bound human PDXP [http://doi.org/10.2210/pdb2CFT/pdb] are used in this manuscript. X-ray crystallographic data of 7,8-DHF-bound murine PDXP generated in this study have been deposited in the PDB and can be accessed under the PDB entry 8QFW [http://doi.org/10.2210/pdb8QFW/pdb]. X-ray crystallographic data of 7,8-DHF-bound human PDXP generated in this study can be accessed under the PDB entries 8S8A (without phosphate) [http://doi.org/10.2210/pdb8S8A/pdb] and 9EM1 (with phosphate) [http://doi.org/10.2210/pdb9EM1/pdb]. The corresponding raw diffraction images have been deposited in the Xtal Raw Data Archive and can be accessed under the XRDA entries 8QFW [https://xrda.pdbj.org/entry/8qfw], 8S8A [https://xrda.pdbj.org/entry/8s8a], and 9EM1 [https://xrda.pdbj.org/entry/9em1].

## AUTHOR CONTRIBUTIONS

Conceptualization, A.G.; validation, E.J.; formal analysis, M.B., C.Z., L.W., S.B., M.N., H.S., E.J., A.G.; investigation, M.B., C.Z., L.W., A.K., K.H., S.B., M.N., E.J.; resources, C.V., M.N., J.P.v.K., H.S.; writing – original draft preparation, A.G. with input from E.J. and H.S.; writing – revised manuscript: A.G. and M.B. with input from H.S. and S.B.; visualization, M.B., C.Z., L.W., S.B., E.J., A.G.; supervision, E.J., A.G.; project administration, A.G.; funding acquisition, A.G.

## SUPPLEMENTARY INFORMATION

**Figure 1 – figure supplement 1.**
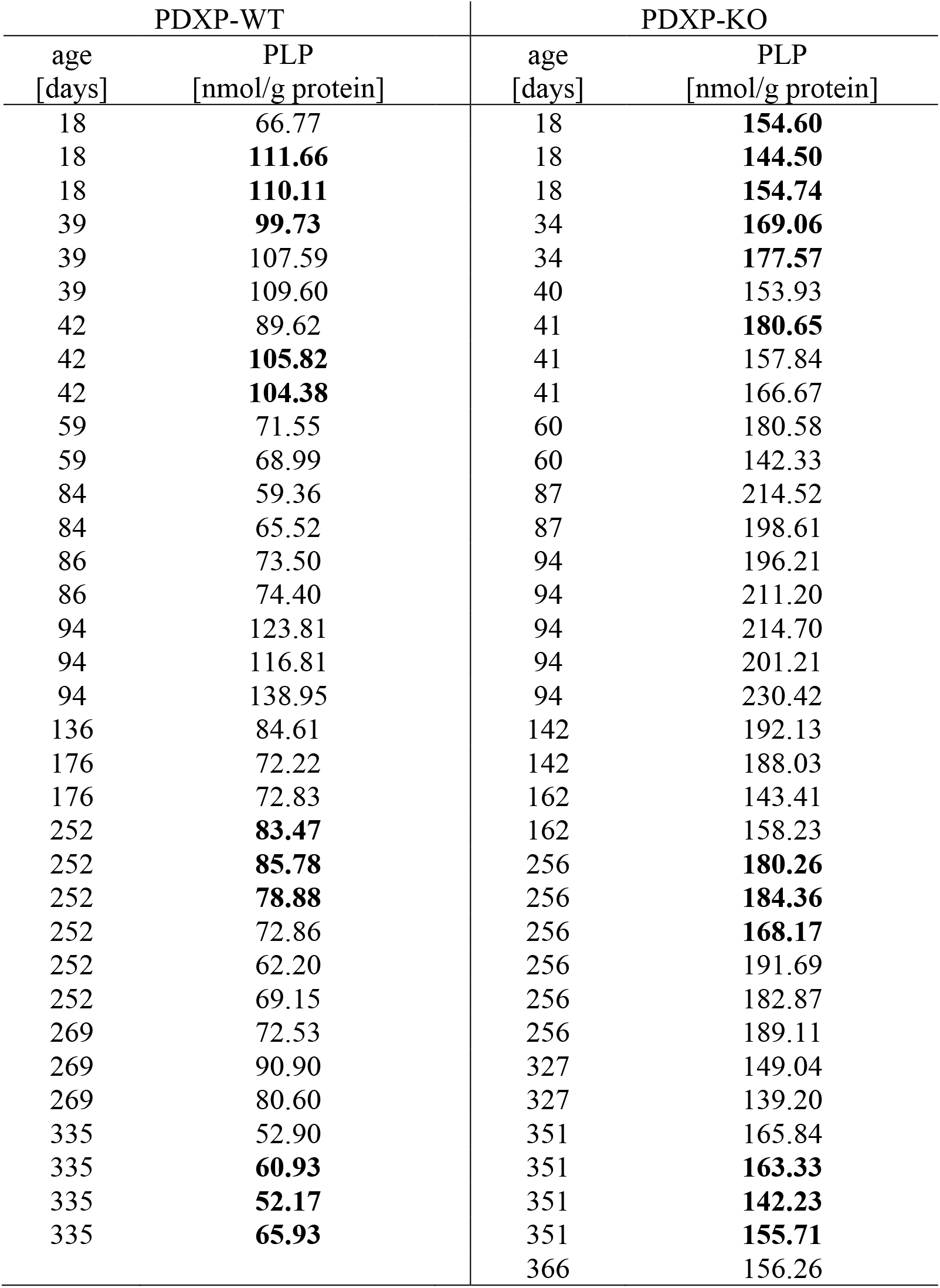
Analysis of total hippocampal PLP levels in PDXP-WT and PDXP-KO mice. Each value represents the result of the PLP determination in an individual hippocampus. Analysis for statistically significant differences between PLP levels in PDXP-WT and PDXP-KO hippocampi (all ages combined; two-tailed, unpaired *t*-test) *p*<0.0001. Bold table entries indicate those hippocampal extracts that were further separated for an analysis of protein-depleted and protein-bound PLP (see Fig. 1c). **Source data** are available for this Table.

**Figure 2 – figure supplement 1.**
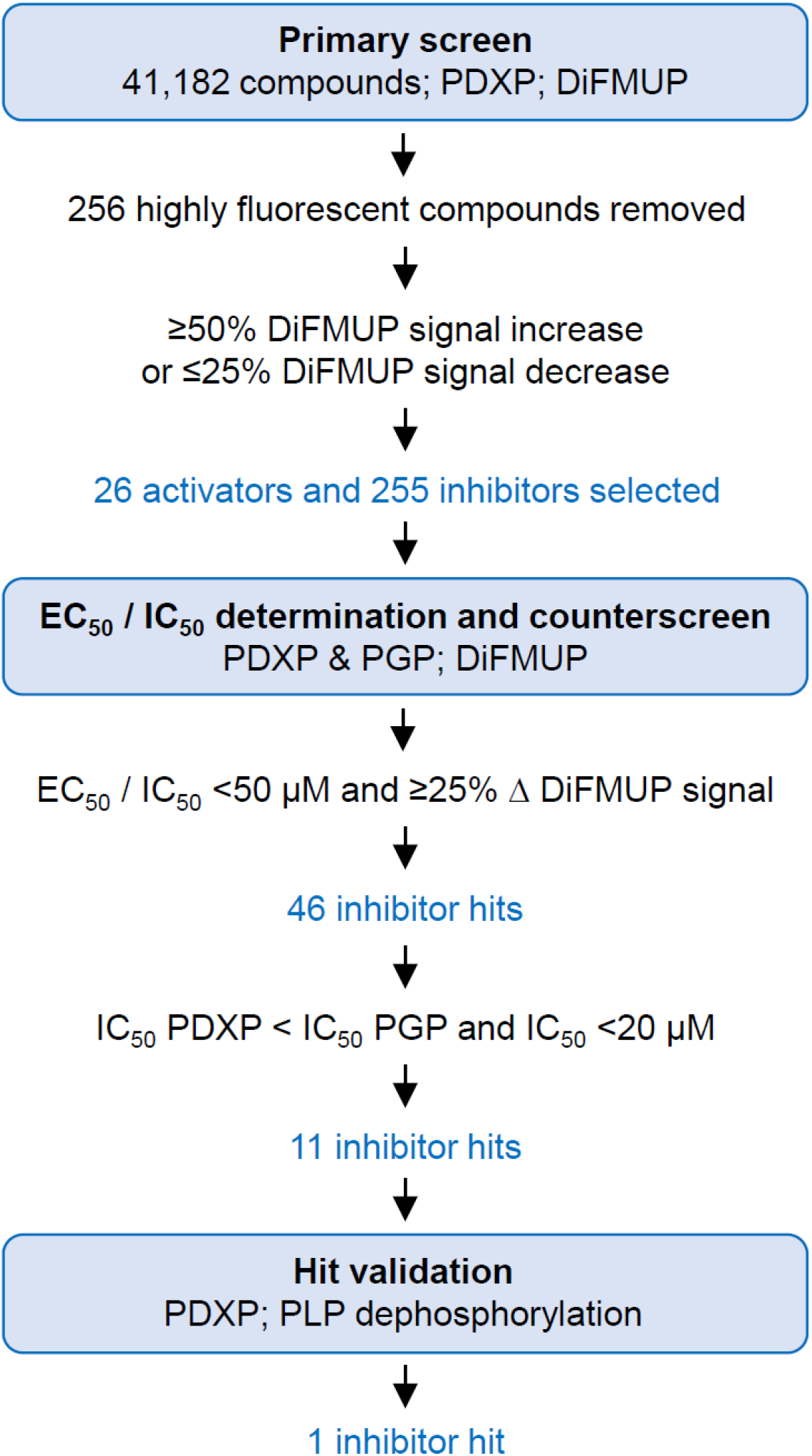
Identification of PDXP inhibitors. A primary screen was conducted using 6,8-difluoro-4-methylumbelliferyl phosphate (DiFMUP) as an artificial substrate. Out of 41,182 screened compounds, 256 compounds were discarded that showed very high autofluorescence (as recognized by elevated fluorescence at the start of the kinetic curve); 26 compounds showed statistically significant PDXP activation, and 255 compounds showed PDXP inhibition (as recognized by an elevated or decreased slope of the kinetic curve, respectively). The average Z’-factor of the screen was 0.75 ± 0.112. These 281 compounds were selected for DiFMUP-based concentration-dependent validation, and the 46 most potent compounds were selected. A counter-screening was conducted in parallel, also in a concentration-dependent fashion, against the PDXP paralog and closest relative phosphoglycolate phosphatase (PGP). The 11 compounds that were inactive against PGP were validated in a secondary assay, using the PDXP substrate pyridoxal 5’-phosphate (PLP). One PDXP inhibitor hit blocked PLP dephosphorylation by ≥50%. **Source data** are available for this Figure.

**Figure 2 – figure supplement 2.**
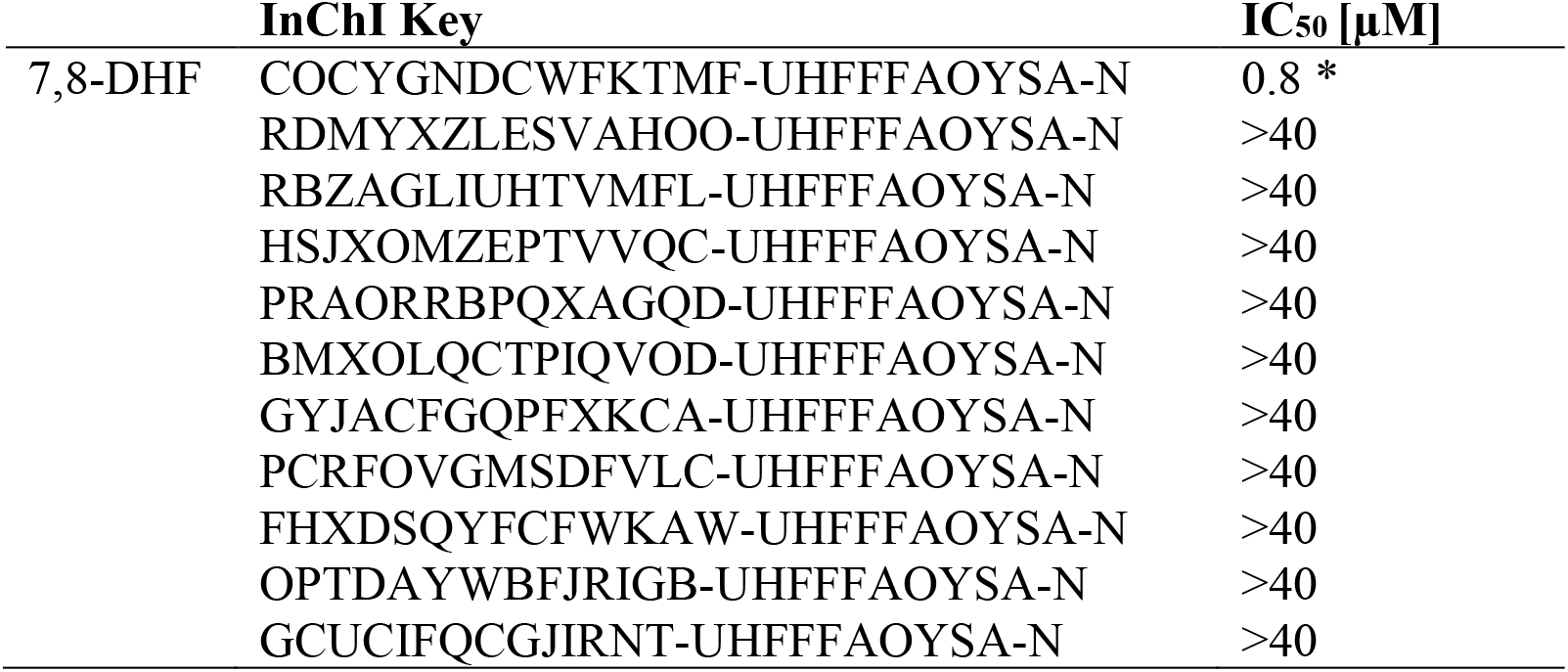
PDXP inhibitor hits. Determination of half-maximal inhibitory constants (IC_50_) of 11 PDXP inhibitory compounds (see InChI Key for chemical substance identification) using purified murine PDXP and pyridoxal 5’-phosphate (PLP) as a substrate. Data marked with an asterisk (*) are results of *n*=3 independent experiments. Because of the limited quantity of most compounds available for these assays, all other data are results of *n*=1 determinations. 7,8-DHF, 7,8-dihydroxyflavone; ∼, approximate IC_50_ value. **Source data** are available for this Table.

**Figure 2 – figure supplement 3.**
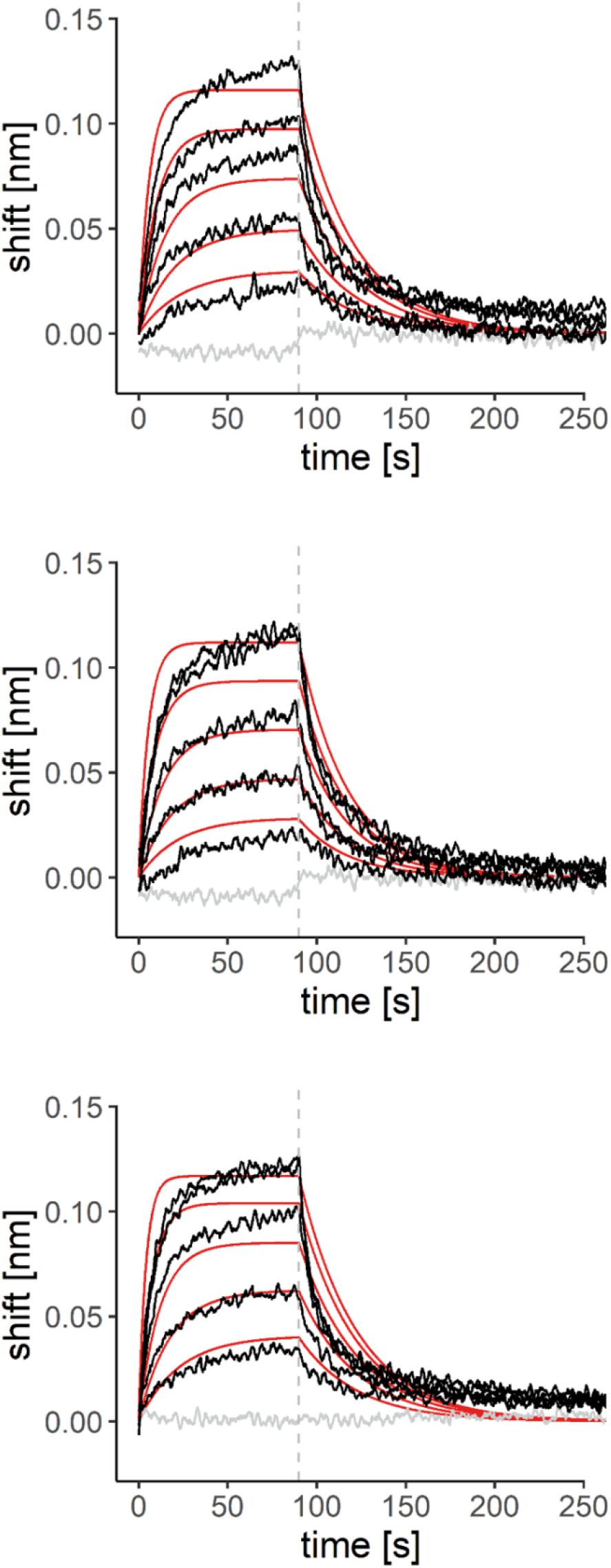
BLI measurements of the interaction of 7,8-DHF with purified murine PDXP. Sensorgrams of three additional experiments overlayed with the global 1:1 binding model (red) and the negative control (gray). The dashed line indicates the start of the dissociation phase. **Source data** are available for this Figure.

**Figure 3 – figure supplement 1.**
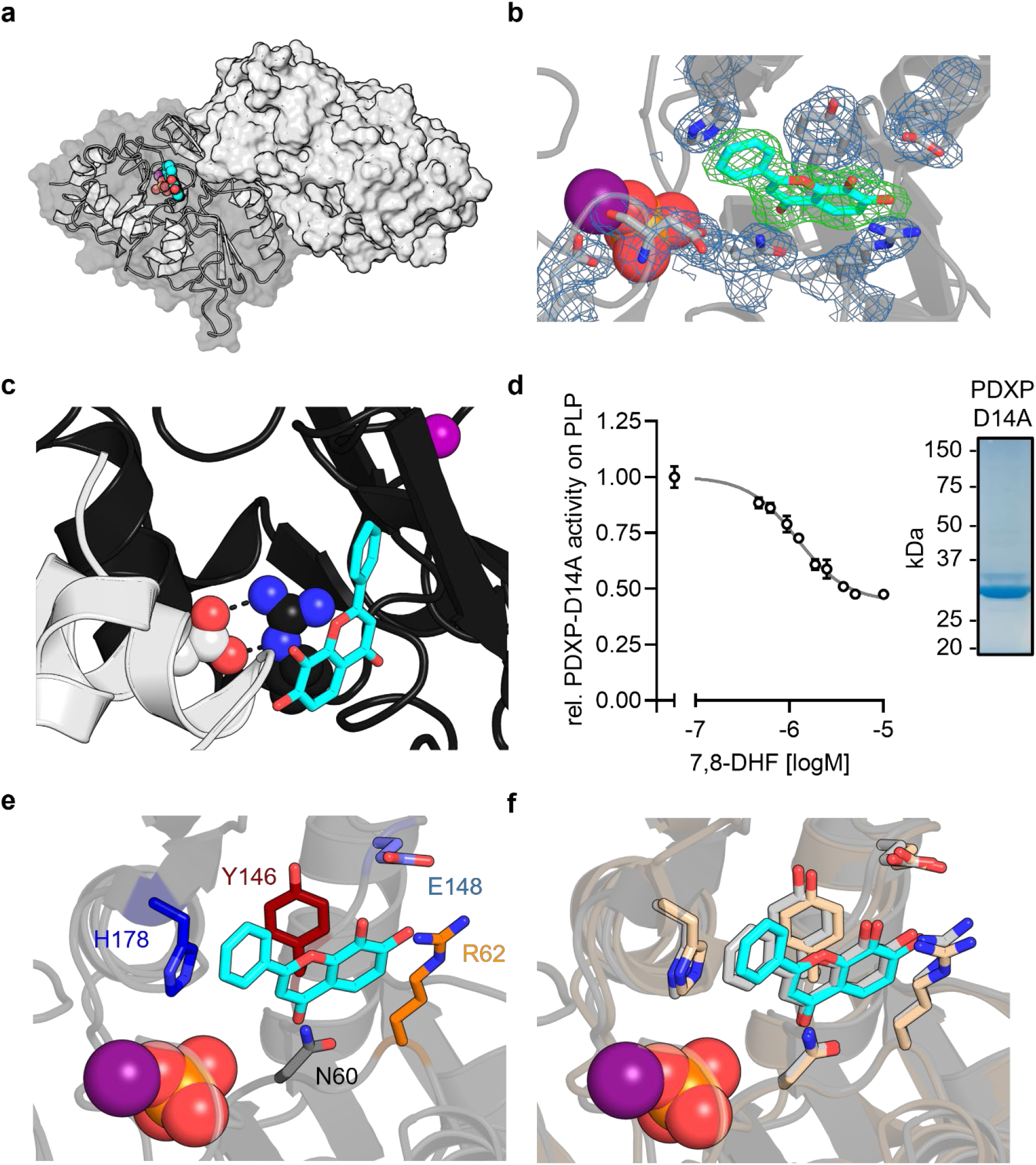
X-ray crystal structures of murine PDXP in complex with 7,8-DHF. (**a**) X-ray crystal structure of the assembled homodimeric, full-length murine PDXP in gray cartoon (protomer A) and surface (protomer B) representation. The model was refined to a resolution of 2.0 Å (PDB code 8QFW). 7,8-DHF is shown in sphere representation with its C-atoms in cyan. The Mg^2+^ ion is shown as deep purple sphere. The phosphate ion is shown as spheres with the phosphorous atom in orange. (**b**) The *2F_o_* − *F_c_* electron density map of the depicted amino acids is contoured at an RMSD of 1.0 in blue mesh and superimposed with the refined model. The *F_o_* − *F_c_* polder electron density map of 7,8-DHF is contoured at an RMSD of 3.0 in green mesh. (**c**) A salt bridge between Arg62 in the B-protomer (in black) and Asp14 of a symmetry-related A-protomer (in gray) blocks the 7,8-DHF binding site in the crystal lattice of murine PDXP. 7,8-DHF (in stick representation with cyan C-atoms) is modeled based on the A-protomer. (**d**) In vitro phosphatase activity of the purified PDXP-D14A variant in the presence of 7,8-DHF. Data are mean values ± S.D. of *n*=3 independent experiments. Apparently missing error bars are hidden by the symbols. The purity of PDXP-D14A is shown in the Coomassie Blue-stained gel on the right. (**e**) Structural details of bound 7,8-DHF and adjacent residues of the active site of mPDXP (in gray cartoon representation). (**f**) Superimposition of the 7,8-DHF binding sites in the phosphate-containing murine PDXP (shown in gray) and human PDXP (in wheat yellow) structures. The corresponding amino acids are shown as gray or wheat yellow-colored sticks. 7,8-DHF bound to murine or human PDXP is shown as sticks colored in gray or cyan, respectively. The position of the Mg^2+^ and phosphate ions is based on human PDXP. **Source data** are available for this Figure.

**Figure 3 – figure supplement 2.**
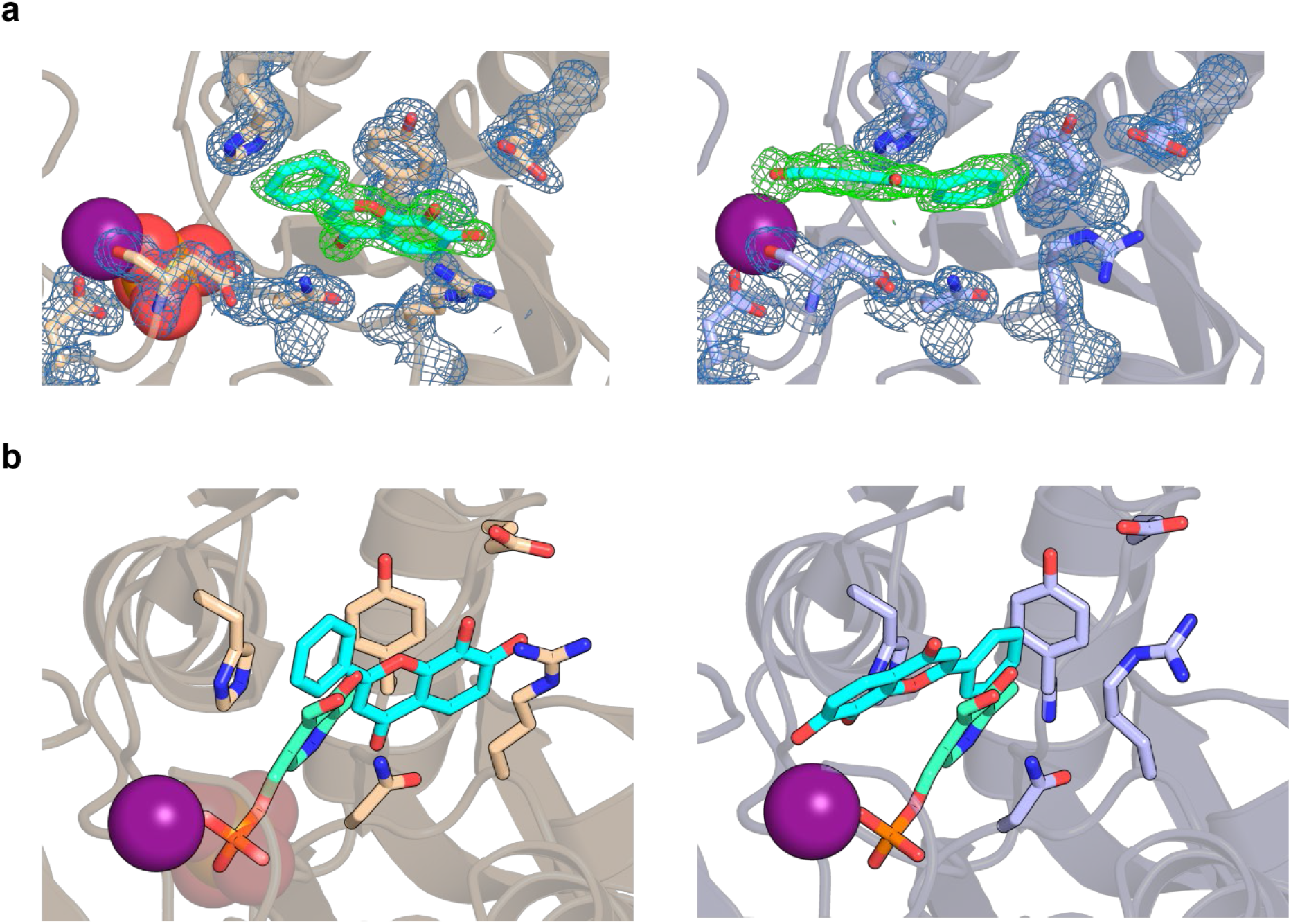
7,8-DHF coordination in PDXP. (**a**) The *2F_o_* − *F_c_* electron density map of the depicted amino acids is contoured at an RMSD of 1.0 in blue mesh and superimposed with the refined model. The *F_o_* − *F_c_* polder electron density map of 7,8-DHF is contoured at an RMSD of 3.0 in green mesh. The Mg^2+^ ion is shown as a deep purple sphere. The phosphate ion is shown as a sphere with the phosphorous atom in orange. *Left panel*, human PDXP with phosphate (cartoon representation in wheat yellow); *right panel*, human PDXP without phosphate (cartoon representation in light blue). (**b**) Comparison of the 7,8-DHF and PLP binding sites in human PDXP with phosphate (wheat yellow, *left panel*), or human PDXP without phosphate (light blue, *right panel*). 7,8-DHF is shown in stick representation with cyan C-atoms. PLP (in stick representation with green C-atoms) was modelled based on a superposition of the human PDXP-PLP complex (PDB code 2CFT).

**Figure 3 – figure supplement 3.**
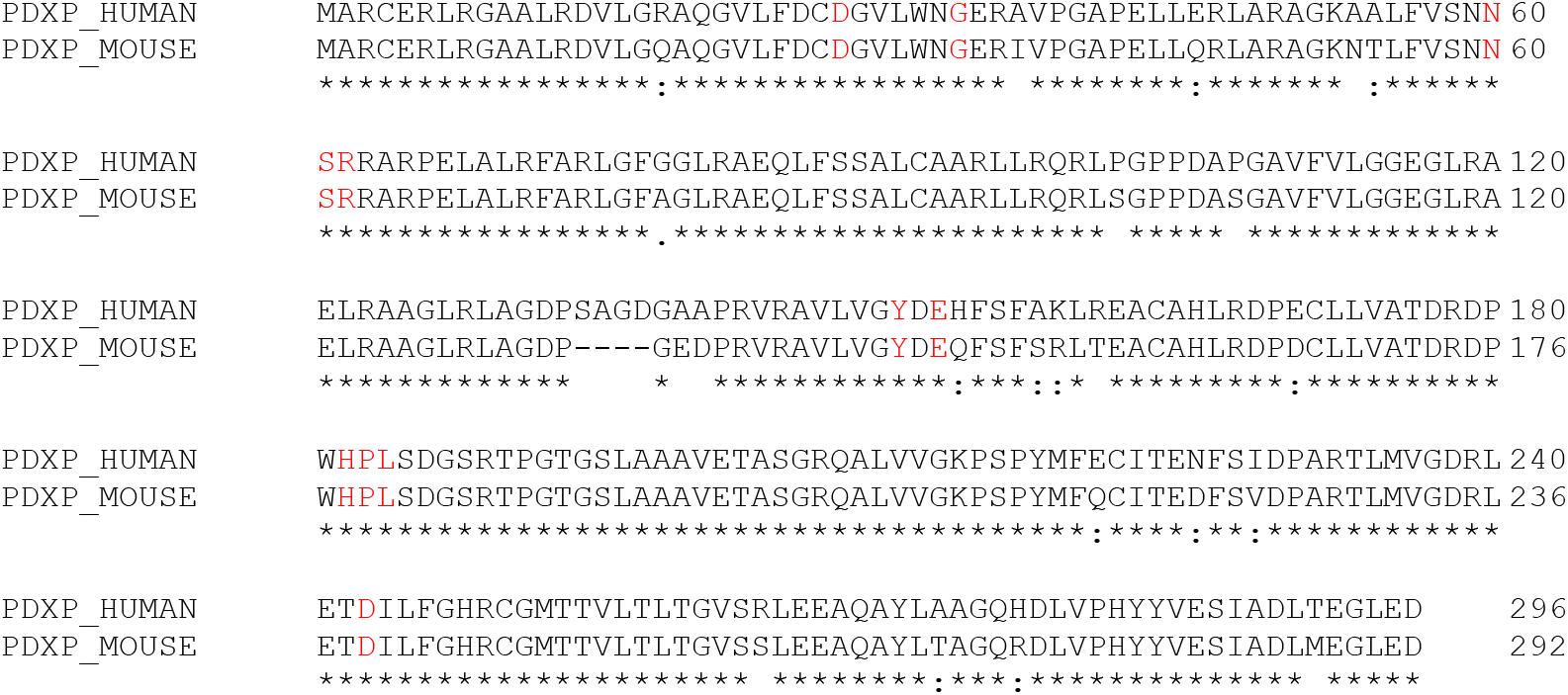
Alignment of human and murine PDXP. Protein sequences of human PDXP (UniProtKB Q96GD0) and murine PDXP (UniProtKB P60487) were aligned with the EMBL-EBI multiple sequence alignment tool Clustal Omega version 1.2.4. PDXP residues found to engage in 7,8-DHF interactions (highlighted in red color) are identical in human and murine PDXP.

**Figure 3 – figure supplement 4.**
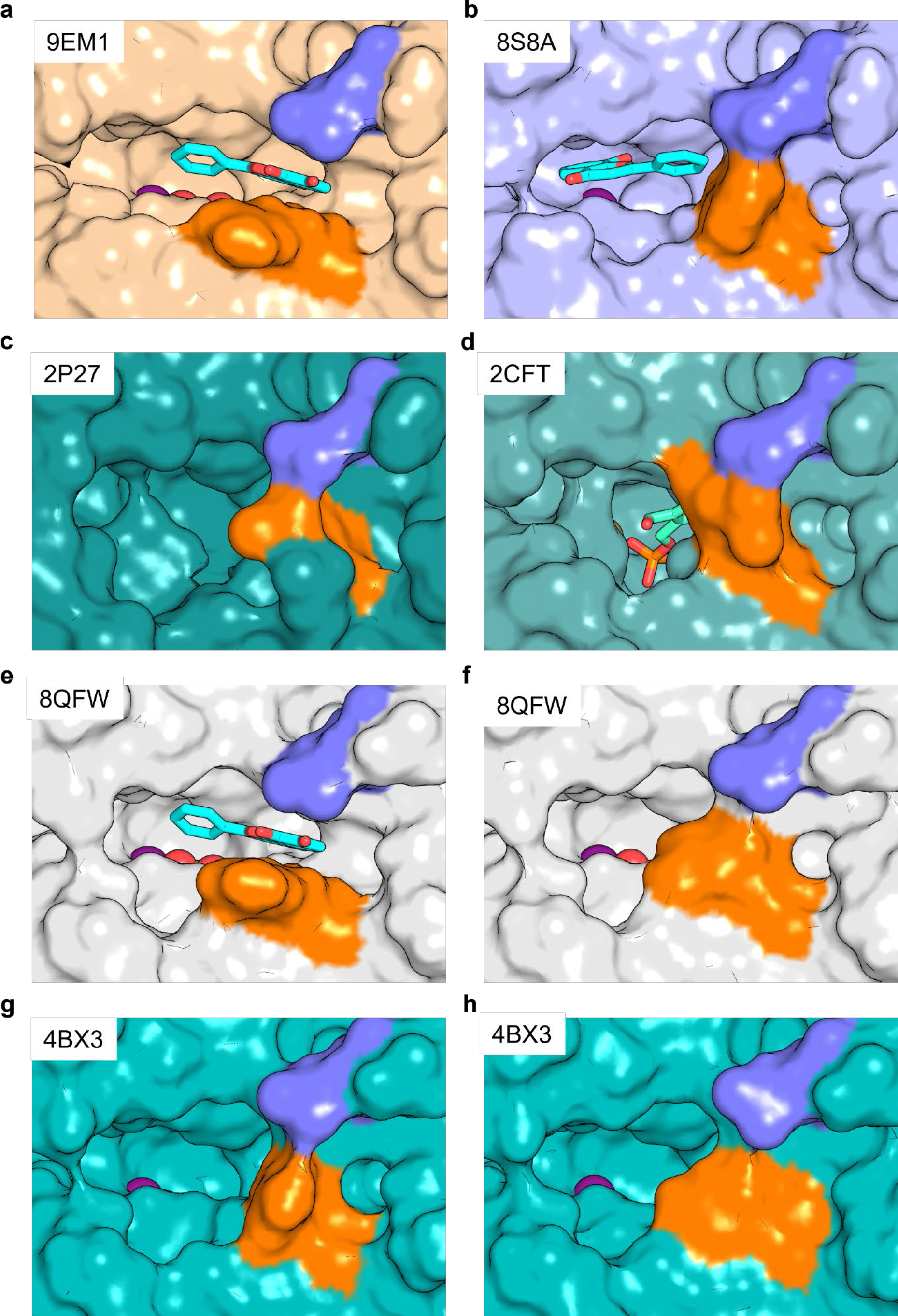
Salt bridge formation between Glu152 (Glu148) and Arg62 gates the active site entrance in PDXP. Shown are views of the active site entrance in (**a**) hPDXP + 7,8-DHF with PO_4_^3-^, (**b**) hPDXP + 7,8-DHF without PO_4_^3-^, (**c**) apo-hPDXP, (**d**) hPDXP + PLP, (**e**) mPDXP + 7,8-DHF with PO_4_^3-^, chain A, (**f**) mPDXP + 7,8-DHF with PO_4_^3-^, chain B (inhibitor-free), (**g**) apo-mPDXP, chain A, (**h**) apo-mPDXP, chain B. The cap domain residue Glu152 in hPDXP (corresponding to Glu148 in mPDXP) is shown in blue, and the core domain residue Arg62 in hPDXP and mPDXP is shown in orange. 7,8-DHF is shown in stick representation with C-atoms in cyan. PLP is shown in stick representation with C-atoms in light green. The phosphate is shown in sphere representation with the phosphorous atom in orange. Mg^2+^ is shown as a deep purple sphere. hPDXP, human PDXP; mPDXP, murine PDXP. The respective PDB entries are indicated.

**Figure 3 – figure supplement 5.**
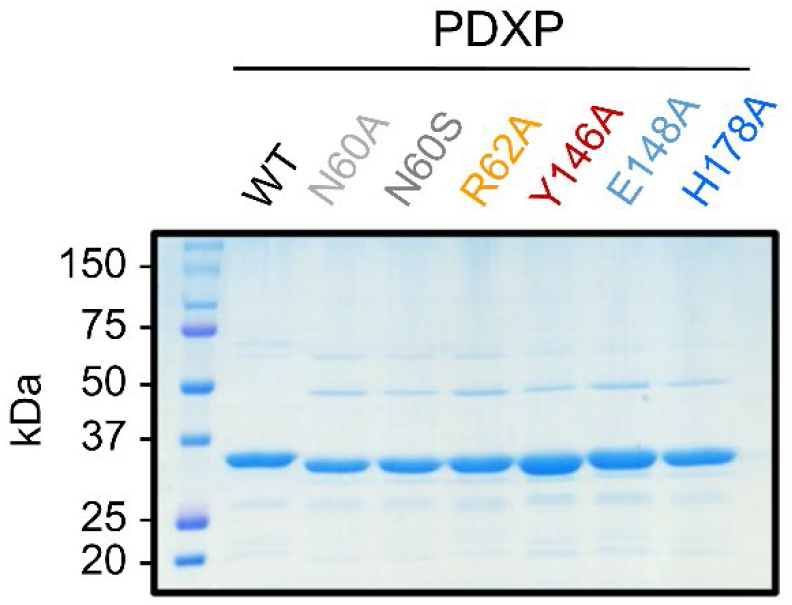
Purity of the employed PDXP and PDXP variants. A Coomassie Blue-stained gel is shown.

## REFERENCES

1. F. G. Bowling, Pyridoxine supply in human development. Semin Cell Dev Biol 22, 611–618 (2011).

2. M. P. Wilson, B. Plecko, P. B. Mills, P. T. Clayton, Disorders affecting vitamin B(6) metabolism. J Inherit Metab Dis 42, 629–646 (2019).

3. M. L. di Salvo, M. K. Safo, R. Contestabile, Biomedical aspects of pyridoxal 5’-phosphate availability. Front Biosci (Elite Ed) 4, 897–913 (2012).

4. E. S. Mitchell, N. Conus, J. Kaput, B vitamin polymorphisms and behavior: evidence of associations with neurodevelopment, depression, schizophrenia, bipolar disorder and cognitive decline. Neurosci Biobehav Rev 47, 307–320 (2014).

5. R. Malouf, J. Grimley Evans, The effect of vitamin B6 on cognition. Cochrane Database Syst Rev 10.1002/14651858.CD004393, CD004393 (2003).

6. C. F. Hughes et al., B-Vitamin Intake and Biomarker Status in Relation to Cognitive Decline in Healthy Older Adults in a 4-Year Follow-Up Study. Nutrients 9 (2017).

7. K. Jannusch et al., A Complex Interplay of Vitamin B1 and B6 Metabolism with Cognition, Brain Structure, and Functional Connectivity in Older Adults. Front Neurosci 11, 596 (2017).

8. H. Xu, S. Wang, F. Gao, C. Li, Vitamin B(6), B(9), and B(12) Intakes and Cognitive Performance in Elders: National Health and Nutrition Examination Survey, 2011-2014. Neuropsychiatr Dis Treat 18, 537–553 (2022).

9. M. F. Elias et al., Homocysteine, folate, and vitamins B6 and B12 blood levels in relation to cognitive performance: the Maine-Syracuse study. Psychosom Med 68, 547–554 (2006).

10. Y. Tomioka et al., Decreased serum pyridoxal levels in schizophrenia: meta-analysis and Mendelian randomization analysis. J Psychiatry Neurosci 43, 194–200 (2018).

11. K. Toriumi et al., Vitamin B6 deficiency hyperactivates the noradrenergic system, leading to social deficits and cognitive impairment. Transl Psychiatry 11, 262 (2021).

12. M. Arai et al., Enhanced carbonyl stress in a subpopulation of schizophrenia. Arch Gen Psychiatry 67, 589–597 (2010).

13. B. D. Paul, Neuroprotective Roles of the Reverse Transsulfuration Pathway in Alzheimer’s Disease. Front Aging Neurosci 13, 659402 (2021).

14. P. M. Ueland, A. McCann, O. Midttun, A. Ulvik, Inflammation, vitamin B6 and related pathways. Mol Aspects Med 53, 10–27 (2017).

15. L. G. Danielski et al., Vitamin B(6) Reduces Neurochemical and Long-Term Cognitive Alterations After Polymicrobial Sepsis: Involvement of the Kynurenine Pathway Modulation. Mol Neurobiol 55, 5255–5268 (2018).

16. Z. Wang, W. Zhu, Y. Xing, J. Jia, Y. Tang, B vitamins and prevention of cognitive decline and incident dementia: a systematic review and meta-analysis. Nutr Rev 80, 931–949 (2022).

17. A. Behrens, E. Graessel, A. Pendergrass, C. Donath, Vitamin B-Can it prevent cognitive decline? A systematic review and meta-analysis. Syst Rev 9, 111 (2020).

18. B. Hassel, A. G. Rogne, S. Hope, Intellectual Disability Associated With Pyridoxine-Responsive Epilepsies: The Need to Protect Cognitive Development. Front Psychiatry 10, 116 (2019).

19. A. W. Rutjes et al., Vitamin and mineral supplementation for maintaining cognitive function in cognitively healthy people in mid and late life. Cochrane Database Syst Rev 12, CD011906 (2018).

20. A. D. Smith, H. Refsum, Homocysteine, B Vitamins, and Cognitive Impairment. Annu Rev Nutr 36, 211–239 (2016).

21. P. S. Aisen et al., High-dose B vitamin supplementation and cognitive decline in Alzheimer disease: a randomized controlled trial. JAMA 300, 1774–1783 (2008).

22. G. Douaud et al., Preventing Alzheimer’s disease-related gray matter atrophy by B-vitamin treatment. Proc Natl Acad Sci U S A 110, 9523–9528 (2013).

23. R. Percudani, A. Peracchi, A genomic overview of pyridoxal-phosphate-dependent enzymes. EMBO Rep 4, 850–854 (2003).

24. A. C. Eliot, J. F. Kirsch, Pyridoxal phosphate enzymes: mechanistic, structural, and evolutionary considerations. Annu Rev Biochem 73, 383–415 (2004).

25. R. Percudani, A. Peracchi, The B6 database: a tool for the description and classification of vitamin B6-dependent enzymatic activities and of the corresponding protein families. BMC Bioinformatics 10, 273 (2009).

26. M. Parra, S. Stahl, H. Hellmann, Vitamin B(6) and Its Role in Cell Metabolism and Physiology. Cells 7 (2018).

27. J. Ciapaite et al., Maintenance of cellular vitamin B(6) levels and mitochondrial oxidative function depend on pyridoxal 5’-phosphate homeostasis protein (PLPHP). J Biol Chem 10.1016/j.jbc.2023.105047, 105047 (2023).

28. A. Fux, S. A. Sieber, Biochemical and Proteomic Studies of Human Pyridoxal 5’-Phosphate-Binding Protein (PLPBP). ACS Chem Biol 15, 254–261 (2020).

29. Y. M. Jang et al., Human pyridoxal phosphatase. Molecular cloning, functional expression, and tissue distribution. J Biol Chem 278, 50040–50046 (2003).

30. E. Jeanclos et al., Improved cognition, mild anxiety-like behavior and decreased motor performance in pyridoxal phosphatase-deficient mice. Biochim Biophys Acta Mol Basis Dis 1865, 193–205 (2019).

31. C. Liu, C. B. Chan, K. Ye, 7,8-dihydroxyflavone, a small molecular TrkB agonist, is useful for treating various BDNF-implicated human disorders. Transl Neurodegener 5, 2 (2016).

32. C. S. Wang, E. T. Kavalali, L. M. Monteggia, BDNF signaling in context: From synaptic regulation to psychiatric disorders. Cell 185, 62–76 (2022).

33. U. Boltaev et al., Multiplex quantitative assays indicate a need for reevaluating reported small-molecule TrkB agonists. Sci Signal 10 (2017).

34. P. Pankiewicz et al., Do Small Molecules Activate the TrkB Receptor in the Same Manner as BDNF? Limitations of Published TrkB Low Molecular Agonists and Screening for Novel TrkB Orthosteric Agonists. Pharmaceuticals (Basel) 14 (2021).

35. D. Todd et al., A monoclonal antibody TrkB receptor agonist as a potential therapeutic for Huntington’s disease. PLoS One 9, e87923 (2014).

36. J. Chen et al., Antioxidant activity of 7,8-dihydroxyflavone provides neuroprotection against glutamate-induced toxicity. Neurosci Lett 499, 181–185 (2011).

37. M. L. Fonda, D. K. Eggers, S. Auerbach, L. Fritsch, Vitamin B-6 metabolism in the brains of young adult and senescent mice. Exp Gerontol 15, 473–479 (1980).

38. A. Gohla, Do metabolic HAD phosphatases moonlight as protein phosphatases? Biochim Biophys Acta Mol Cell Res 1866, 153–166 (2019).

39. A. Seifried et al., Evolutionary and structural analyses of mammalian haloacid dehalogenase-type phosphatases AUM and chronophin provide insight into the basis of their different substrate specificities. J Biol Chem 289, 3416–3431 (2014).

40. E. Jeanclos et al., Glycolytic flux control by drugging phosphoglycolate phosphatase. Nat Commun 13, 6845 (2022).

41. M. Congreve, R. Carr, C. Murray, H. Jhoti, A ‘rule of three’ for fragment-based lead discovery? Drug Discov Today 8, 876–877 (2003).

42. J. O’Connell et al., Small molecules that inhibit TNF signalling by stabilising an asymmetric form of the trimer. Nat Commun 10, 5795 (2019).

43. A. Seifried, J. Schultz, A. Gohla, Human HAD phosphatases: structure, mechanism, and roles in health and disease. Febs j 280, 549–571 (2013).

44. A. M. Burroughs, K. N. Allen, D. Dunaway-Mariano, L. Aravind, Evolutionary genomics of the HAD superfamily: understanding the structural adaptations and catalytic diversity in a superfamily of phosphoesterases and allied enzymes. J Mol Biol 361, 1003–1034 (2006).

45. H. Huang et al., Panoramic view of a superfamily of phosphatases through substrate profiling. Proc Natl Acad Sci U S A 112, E1974–1983 (2015).

46. C. Proenca et al., Inhibition of protein tyrosine phosphatase 1B by flavonoids: A structure - activity relationship study. Food Chem Toxicol 111, 474–481 (2018).

47. V. Bianchi, J. Spychala, Mammalian 5’-nucleotidases. J Biol Chem 278, 46195–46198 (2003).

48. C. Kestler et al., Chronophin dimerization is required for proper positioning of its substrate specificity loop. J Biol Chem 289, 3094–3103 (2014).

49. G. Knobloch et al., Synthesis of hydrolysis-resistant pyridoxal 5’-phosphate analogs and their biochemical and X-ray crystallographic characterization with the pyridoxal phosphatase chronophin. Bioorg Med Chem 23, 2819–2827 (2015).

50. Z. Zhang et al., 7,8-dihydroxyflavone prevents synaptic loss and memory deficits in a mouse model of Alzheimer’s disease. Neuropsychopharmacology 39, 638–650 (2014).

51. L. Devi, M. Ohno, 7,8-dihydroxyflavone, a small-molecule TrkB agonist, reverses memory deficits and BACE1 elevation in a mouse model of Alzheimer’s disease. Neuropsychopharmacology 37, 434–444 (2012).

52. E. Bollen et al., 7,8-Dihydroxyflavone improves memory consolidation processes in rats and mice. Behav Brain Res 257, 8–12 (2013).

53. N. A. Castello et al., 7,8-Dihydroxyflavone, a small molecule TrkB agonist, improves spatial memory and increases thin spine density in a mouse model of Alzheimer disease-like neuronal loss. PLoS One 9, e91453 (2014).

54. N. Aytan et al., Protective effects of 7,8-dihydroxyflavone on neuropathological and neurochemical changes in a mouse model of Alzheimer’s disease. Eur J Pharmacol 828, 9–17 (2018).

55. L. Gao et al., TrkB activation by 7, 8-dihydroxyflavone increases synapse AMPA subunits and ameliorates spatial memory deficits in a mouse model of Alzheimer’s disease. J Neurochem 136, 620–636 (2016).

56. A. Akhtar, J. Dhaliwal, S. P. Sah, 7,8-Dihydroxyflavone improves cognitive functions in ICV-STZ rat model of sporadic Alzheimer’s disease by reversing oxidative stress, mitochondrial dysfunction, and insulin resistance. Psychopharmacology (Berl) 238, 1991–2009 (2021).

57. Y. H. Hsiao, H. C. Hung, S. H. Chen, P. W. Gean, Social interaction rescues memory deficit in an animal model of Alzheimer’s disease by increasing BDNF-dependent hippocampal neurogenesis. J Neurosci 34, 16207–16219 (2014).

58. A. Blugeot et al., Vulnerability to depression: from brain neuroplasticity to identification of biomarkers. J Neurosci 31, 12889–12899 (2011).

59. J. C. Zhang et al., Antidepressant effects of TrkB ligands on depression-like behavior and dendritic changes in mice after inflammation. Int J Neuropsychopharmacol 18 (2014).

60. W. Yao et al., Role of Keap1-Nrf2 signaling in depression and dietary intake of glucoraphanin confers stress resilience in mice. Sci Rep 6, 30659 (2016).

61. M. W. Zhang, S. F. Zhang, Z. H. Li, F. Han, 7,8-Dihydroxyflavone reverses the depressive symptoms in mouse chronic mild stress. Neurosci Lett 635, 33–38 (2016).

62. Y. Li et al., Inflammation-activated C/EBPbeta mediates high-fat diet-induced depression-like behaviors in mice. Front Mol Neurosci 15, 1068164 (2022).

63. N. Amin et al., Optimized integration of fluoxetine and 7, 8-dihydroxyflavone as an efficient therapy for reversing depressive-like behavior in mice during the perimenopausal period. Prog Neuropsychopharmacol Biol Psychiatry 101, 109939 (2020).

64. E. J. Jaehne et al., TrkB agonist 7,8-dihydroxyflavone reverses an induced prepulse inhibition deficit selectively in maternal immune activation offspring: implications for schizophrenia. Behav Pharmacol 32, 404–412 (2021).

65. M. Han et al., Intake of 7,8-Dihydroxyflavone During Juvenile and Adolescent Stages Prevents Onset of Psychosis in Adult Offspring After Maternal Immune Activation. Sci Rep 6, 36087 (2016).

66. Y. J. Yang et al., Small-molecule TrkB agonist 7,8-dihydroxyflavone reverses cognitive and synaptic plasticity deficits in a rat model of schizophrenia. Pharmacol Biochem Behav 122, 30–36 (2014).

67. M. Han, J. C. Zhang, K. Hashimoto, Increased Levels of C1q in the Prefrontal Cortex of Adult Offspring after Maternal Immune Activation: Prevention by 7,8-Dihydroxyflavone. Clin Psychopharmacol Neurosci 15, 64–67 (2017).

68. Q. Ren et al., Effects of TrkB agonist 7,8-dihydroxyflavone on sensory gating deficits in mice after administration of methamphetamine. Pharmacol Biochem Behav 106, 124–127 (2013).

69. C. Becker et al., Predicting and treating stress-induced vulnerability to epilepsy and depression. Ann Neurol 78, 128–136 (2015).

70. A. Guarino et al., Low-dose 7,8-Dihydroxyflavone Administration After Status Epilepticus Prevents Epilepsy Development. Neurotherapeutics 19, 1951–1965 (2022).

71. R. A. Johnson et al., 7,8-dihydroxyflavone exhibits therapeutic efficacy in a mouse model of Rett syndrome. J Appl Physiol(1985) 112, 704–710 (2012).

72. M. S. Kang et al., Autism-like behavior caused by deletion of vaccinia-related kinase 3 is improved by TrkB stimulation. J Exp Med 214, 2947–2966 (2017).

73. Y. Lee, P. L. Han, Early-Life Stress in D2 Heterozygous Mice Promotes Autistic-like Behaviors through the Downregulation of the BDNF-TrkB Pathway in the Dorsal Striatum. Exp Neurobiol 28, 337–351 (2019).

74. Y. S. Chen et al., Early 7,8-Dihydroxyflavone Administration Ameliorates Synaptic and Behavioral Deficits in the Young FXS Animal Model by Acting on BDNF-TrkB Pathway. Mol Neurobiol 60, 2539–2552 (2023).

75. M. Majewski, A. Kozlowska, M. Thoene, E. Lepiarczyk, W. J. Grzegorzewski, Overview of the role of vitamins and minerals on the kynurenine pathway in health and disease. J Physiol Pharmacol 67, 3–19 (2016).

76. M. A. Sorolla et al., Impaired PLP-dependent metabolism in brain samples from Huntington disease patients and transgenic R6/1 mice. Metab Brain Dis 31, 579–586 (2016).

77. S. W. Jang et al., A selective TrkB agonist with potent neurotrophic activities by 7,8-dihydroxyflavone. Proc Natl Acad Sci U S A 107, 2687–2692 (2010).

78. K. Kobayashi, H. Suzuki, Synapse-selective rapid potentiation of hippocampal synaptic transmission by 7,8-dihydroxyflavone. Neuropsychopharmacol Rep 38, 197–203 (2018).

79. D. Talwar et al., Optimisation and validation of a sensitive high-performance liquid chromatography assay for routine measurement of pyridoxal 5-phosphate in human plasma and red cells using pre-column semicarbazide derivatisation. Journal of chromatography. B, Analytical technologies in the biomedical and life sciences 792, 333–343 (2003).

80. F. W. Studier, Protein production by auto-induction in high density shaking cultures. Protein Expr Purif 41, 207–234 (2005).

81. C. A. Wartchow et al., Biosensor-based small molecule fragment screening with biolayer interferometry. J Comput Aided Mol Des 25, 669–676 (2011).

82. W. Kabsch, Xds. Acta crystallographica. Section D, Biological crystallography 66, 125–132 (2010).

83. P. R. Evans, G. N. Murshudov, How good are my data and what is the resolution? Acta crystallographica. Section D, Biological crystallography 69, 1204–1214 (2013).

84. M. D. Winn et al., Overview of the CCP4 suite and current developments. Acta crystallographica. Section D, Biological crystallography 67, 235–242 (2011).

85. A. J. McCoy et al., Phaser crystallographic software. J Appl Crystallogr 40, 658–674 (2007).

86. P. D. Adams et al., PHENIX: a comprehensive Python-based system for macromolecular structure solution. Acta crystallographica. Section D, Biological crystallography 66, 213–221 (2010).

87. P. Emsley, B. Lohkamp, W. G. Scott, K. Cowtan, Features and development of Coot. Acta crystallographica. Section D, Biological crystallography 66, 486–501 (2010).

88. Schrodinger, LLC (2021) The PyMOL Molecular Graphics System, Version 2.5.1.

